# CRISPR/Cas9-based repeat depletion for the high-throughput genotyping of complex plant genomes

**DOI:** 10.1101/2022.11.22.517518

**Authors:** Marzia Rossato, Luca Marcolungo, Luca De Antoni, Giulia Lopatriello, Elisa Bellucci, Gaia Cortinovis, Giulia Frascarelli, Laura Nanni, Elena Bitocchi, Valerio Di Vittori, Leonardo Vincenzi, Filippo Lucchini, Kirstin E. Bett, Larissa Ramsay, David James Konkin, Massimo Delledonne, Roberto Papa

## Abstract

High-throughput genotyping enables the large-scale analysis of genetic diversity in population genomics and genomewide association studies that combine the genotypic and phenotypic characterization of large collections of accessions. Sequencing-based approaches for genotyping are progressively replacing traditional genotyping methods due to the lower ascertainment bias. However, genome-wide genotyping based on sequencing becomes expensive in species with large genomes and a high proportion of repetitive DNA. Here we describe the use of CRISPR/Cas9 technology to deplete repetitive elements in the 3.76-Gb genome of lentil (*Lens culinaris*), 84% consisting of repeats, thus concentrating the sequencing data on coding and regulatory regions (single-copy regions). We designed a custom set of 566,766 gRNAs targeting 2.9 Gbp of repeats and excluding repetitive regions overlapping annotated genes and putative regulatory elements based on ATAC-Seq data. The novel depletion method removed ∼40% of reads mapping to repeats, increasing those mapping to single-copy regions by ∼2.6-fold. When analyzing 25 million fragments, this repeat-to-single-copy shift in the sequencing data increased the number of genotyped bases of ∼10-fold compared to non-depleted libraries. In the same condition, we were also able to identify ∼12-fold more genetic variants in the single-copy regions and increased the genotyping accuracy by rescuing thousands of heterozygous variants that otherwise would be missed due to low coverage. The method performed similarly regardless of the multiplexing level, type of library or genotypes, including different cultivars and a closely-related species (*L. orientalis*). Our results demonstrated that CRISPR/Cas9-driven repeat depletion focuses sequencing data on meaningful genomic regions, thus improving high-density and genome-wide genotyping in large and repetitive genomes.

## INTRODUCTION

The efficient and accurate determination of genotypes is necessary for large-scale projects investigating the genetic composition of germplasm collections representing wild and domesticated species and inbred lines. One example is the EU H2020 project INCREASE (www.pulsesincrease.eu) (Bellucci et al. 2021), which focuses on four legume staples: chickpea, common bean, lentil and lupin. Such projects depend on large cohorts of individuals to enable the comparative analysis of samples with sufficient statistical power. Cost-effective high-throughput genotyping methods are therefore needed to increase the number of samples that can be processed in an economically feasible manner (Bellucci et al. 2021). This can only be achieved by reducing the fraction of each individual genome that is sequenced while ensuring that the same homologous regions are examined in each individual (Peterson et al. 2012).

High-throughput low-cost genotyping has largely been achieved by the analysis of single-nucleotide polymorphisms on microarray-based platforms (SNP arrays). These allow up to several thousand SNPs to be tested simultaneously (Pavan et al. 2020). This approach considers a predefined set of markers, resulting in fixed costs per individual regardless of the genome size and fraction of repetitive DNA. However, analysis is restricted to known SNPs that are frequent in the population, while rare and unknown SNPs are ignored. This is a drawback when analyzing diverse landraces and distant wild relatives, as required in the germplasm characterization projects mentioned above (Lachance and Tishkoff 2013).

More recently, next generation sequencing (NGS) has provided an opportunity to discover genome-wide variants in a less biased manner. Sequencing-based approaches for genotyping involves low coverage (5–10x) whole-genome sequencing (lcWGS), allowing the characterization of several million variants (Tanaka et al. 2021; Friel et al. 2021). To reduce costs enough to make WGS affordable even in large germplasm collections, very low coverage (0.5–2x) WGS (ultra-lcWGS) can be combined with imputation to infer positions that are not sequenced or genotyped (Deng et al. 2022; Zan et al. 2019; Wang et al. 2016). Alternatively, sequencing costs are often minimized by reduced-representation sequencing, which comprises methods such as genotyping by sequencing (GBS) (Elshire et al. 2011), restriction siteassociated DNA sequencing (RAD-Seq) (Davey et al. 2011; Baird et al. 2008) and double-digest RAD-Seq (ddRAD-seq) (Truong et al. 2012; Peterson et al. 2012). These methods concentrate sequencing data on regions adjacent to restriction sites by exploiting the specificity of restriction endonucleases. Reduced-representation sequencing is suitable for large cohorts, but provides only low-resolution data, with a small fraction of analysed and genotyped bases (Pavan et al. 2020) that may not provide sufficient marker density and depth to confidently identify variants under selection in large genomes (Guerra-García et al. 2021). The resolution can be increased without significantly greater costs in sample prep by using the Twist 96-Plex Library Prep Kit (formerly iGenomX Riptide Kit) to generate multiplexed libraries, allowing 96–960 samples to be processed simultaneously and resulting in the non-random sampling of millions of genomic positions (Siddique et al. 2019).

Despite the advantages of sequencing-based genotyping over SNP arrays, one common disadvantage is that sequencing methods generally do not distinguish between repetitive (low-complexity) and single-copy (high-complexity) regions, the latter comprising coding and regulatory regions that are the main targets of natural selection and thus the focus of most genotyping projects. In contrast, low-complexity regions of plant genomes mainly comprise transposable elements, simple sequence repeats and tandem repeats. Transposable elements play a key role in genome evolution, but the analysis of such regions is technically challenging and largely uninformative in genotyping studies, unless dedicated analysis workflows are applied (Yan et al. 2022). Mapping reads to transposable/repetitive elements can result in lowquality alignments that hinder the calling of accurate genotypes, which is a consistent challenge particularly for those plant species with large genomes, where repetitive elements account for up 90% of the total DNA. This includes many domesticated crops such as corn (*Zea mays*), wheat (*Triticum* spp.), lentil (*Lens culinaris*) and onion (*Allium cepa*) (Feuillet et al. 2011). One strategy to address this issue is whole exome sequencing (WES), which selects coding regions for preferential sequencing (Hodges et al. 2007) as shown in lentil, wheat and barley (Ogutcen et al. 2018; He et al. 2019). However, WES only focuses on coding sequences and thus overlooks regulatory elements, which are equally important as sources of genetic diversity (Ricci et al. 2019; Wang et al. 2019; Tian et al. 2020).

Ideally, lcWGS could be focused on the most complex parts of the genome, avoiding wasted effort on the sequencing of repetitive elements. This could be achieved by using enzymes that enable target enrichment by depleting unwanted sequences from NGS libraries. For example, the duplex-specific nuclease (DSN) selectively digests double-stranded DNA molecules, and can be used to eliminate highly abundant sequences in a controlled denaturation-reassociation reaction (Zhulidov et al. 2004). This method has been used in RNA-Seq analysis to remove abundant transcripts (Zhao et al. 2014; Miller et al. 2013) and, just occasionally, also to delete repetitive elements in DNA-Seq libraries generated from plant genomes (Ichida and Abe 2019; Matvienko et al. 2013). However, (Matvienko et al. 2013)DSN can also remove informative repetitive elements, such as the coding sequences of abundant gene families, which are particularly relevant in polyploid plants arising from whole genome duplication events (Matvienko et al. 2013). More recently, the CRISPR/Cas9 (clustered regularly interspaced short palindromic repeats and CRISPR-associated nuclease 9) system has been used for the selective depletion of unwanted genome fractions from sequencing libraries (Gu et al. 2016). The Cas9 enzyme can be programmed to cut library fragments by designing specific guide-RNA sequences targeting the unwanted sequences. Subsequently, only intact fragments -retaining adapters at both ends-can be effectively amplified by PCR and generate productive clusters on a sequencing flow-cell. The DASH approach (depletion of abundant sequences by hybridization) involved the use of Cas9 to exclude ribosomal RNA (rRNA) sequences from RNA-Seq libraries and to remove DNA from common pathogens in order to detect rare pathogens in metagenomic samples (Gu et al. 2016). A similar technology has been recently commercialized under the name “CRISPRclean” by JumpCode Genomics (JumpCode Genomics 2021).

Here we determined whether CRISPRclean technology could be used to deplete the repetitive elements in libraries prepared from the 3.76-Gbp genome of lentil (*L. culinaris*), 84% of which is repetitive DNA. CRISPRclean technology was combined with Twist multiplexing libraries and we evaluated its performance, focusing on the technical features required for genotyping. Our results will facilitate the large-scale genomic analysis of lentil as well as other plant species with large and highly-repetitive genomes.

## RESULTS

### Depletion of *L. culinaris* repetitive DNA using CRISPR/Cas9

We designed a custom set of gRNAs to deplete the repetitive DNA content of the *L. culinaris* CDC Redberry genome (Ramsay et al. 2021), targeting transposable elements (totaling 3.1 Gbp of sequence across the whole genome, corresponding to 82.5% of its size), simple sequence repeats (58 Mbp, 1.5%) and tandem repeats (13 Mbp, 0.4%) in the nuclear genome, as well as the entire mitochondrial genome (mtDNA, 489 kbp) and chloroplast genome (cpDNA, 118 kbp) (**Supplemental Table S1**). We excluded repetitive DNA that overlapped with functional regions such as annotated genes (185 Mbp, 5%) and putative regulatory regions identified using ATAC-Seq data (78 Mbp, 2%) (**Supplemental Table S1**). All nuclear DNA outside the gRNA target regions is hereafter defined as single-copy. The final design comprised 566,766 gRNAs with at least 25 recognition sites, potentially targeting 2.9 Gbp (77%) of the *L. culinaris* nuclear genome and 93.5% of its repetitive regions when using a sequencing library with 500-bp inserts (**Supplemental Table S2** and **Supplemental File S1**). An additional 2,366 gRNAs targeted the mitochondrial and chloroplast genomes (**Supplemental Table S2**). The gRNAs were assigned to 11 pools based on their cutting frequency in the *L. culinaris* genome (**Supplemental Table S2**).

The custom gRNAs were tested on three Twist 8-plex libraries, allowing the reproducible sampling of the same genomic regions by random priming during first-strand DNA synthesis. Each 8-plex library comprised replicates of three distinct *L. culinaris* samples (cv. Castelluccio) (**Supplemental Table S3**). Cas9/gRNAs ribonucleoprotein (RNP) complexes (1:2.5 protein/gRNA ratio) were generated, and gRNAs with more target sites in the genome were used at higher relative concentrations in the final reaction (**Supplemental Table S2**). Depletion reactions in the presence of RNP complexes were conducted either using all gRNAs simultaneously or by splitting the gRNA pools into three groups based on cutting frequency (**Supplemental Table S2**) and using the groups sequentially, starting with the lowest cutting frequency. Depleted and non-depleted libraries were sequenced, generating 91 million fragments on average (**Supplemental Table S4**). The sequencing data were normalized at ∼50 million fragments per library in order to compare the proportion of reads mapping on repetitive and single-copy regions of the nuclear genome and on the organelle genomes (**Figure 1**). The number of reads mapping to repetitive regions (total repeats, nuclear repeats, mtDNA and cpDNA) was significantly lower in the depleted libraries compared to the non-depleted libraries, with the sequential depletion strategy using three gRNA groups performing best and depleting 37.7% of the repetitive DNA (**Figure 1A**). The results were similar when considering only the nuclear repetitive regions (37.2% depletion) (**Figure 1B**). A small fraction of total reads mapped to the organelle genomes (∼1%). Both depletion strategies were similarly effective in the chloroplast genome, resulting in ∼78.5% depletion (**Figure 1C**). In contrast, no significant depletion was observed in the mitochondrial genome (**Figure 1D**). In parallel, the sequential depletion strategy achieved a 130% increase in the number of reads mapping to singlecopy regions, from 17.5 to 40.4 million (**Figure 1E**). Given that the concentration of Cas9 RNPs influences the cutting efficiency (Gu et al. 2016), we repeated the sequential depletion strategy using double amount of Cas9 and gRNAs. This modified the read distribution further, achieving 41.2% depletion of nuclear repeats and a 160% increase in reads mapped to single-copy regions. The sequential depletion strategy with double RNPs was therefore the most efficient, and was applied in all subsequent experiments. Overall, our results demonstrated that the custom gRNA set and Cas9 effectively targeted fragments containing repetitive DNA sequences and depleted them in the resulting sequencing libraries. **Figure 2A-B** shows the alignments of reads at two representative genomic regions, confirming that less sequencing data was assigned to regions of repetitive DNA and more reads were mapped to single-copy parts of the genome ().

**Figure 1.**
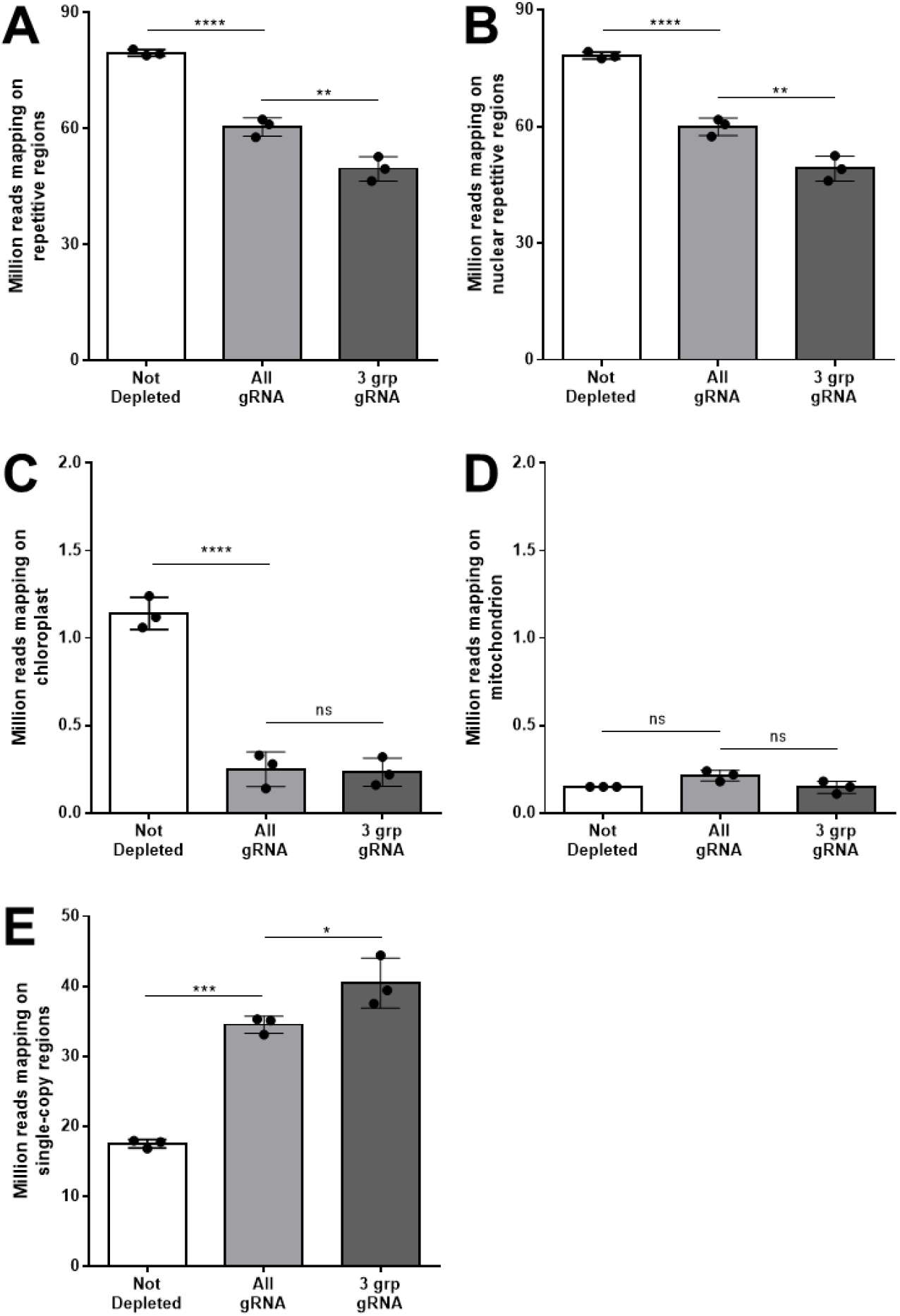
Distribution of mapped reads after CRISPR/Cas9-mediated repeat depletion. Libraries of *L. culinaris* cv. Castelluccio DNA (8-plex) were depleted using the custom gRNA set and Cas9. The gRNAs were used simultaneously (All) or were split into three groups that were used sequentially in order of increasing cutting frequency (3 grp). Bar graphs show the number (in millions) of reads mapping to repetitive DNA (**A**), to nuclear repetitive DNA (**B**), to the chloroplast genome (**C**), to the mitochondrial genome (**D**), and to the single-copy regions of the nuclear genome (**E**). Data are means ± SD (n = 3 for each condition; *p-adj < 0.05, **p-adj < 0.01, ****p-adj < 0.0001; one-way ANOVA plus Tukey’s multiple comparisons test; ns, not significant).

**Figure 2.**
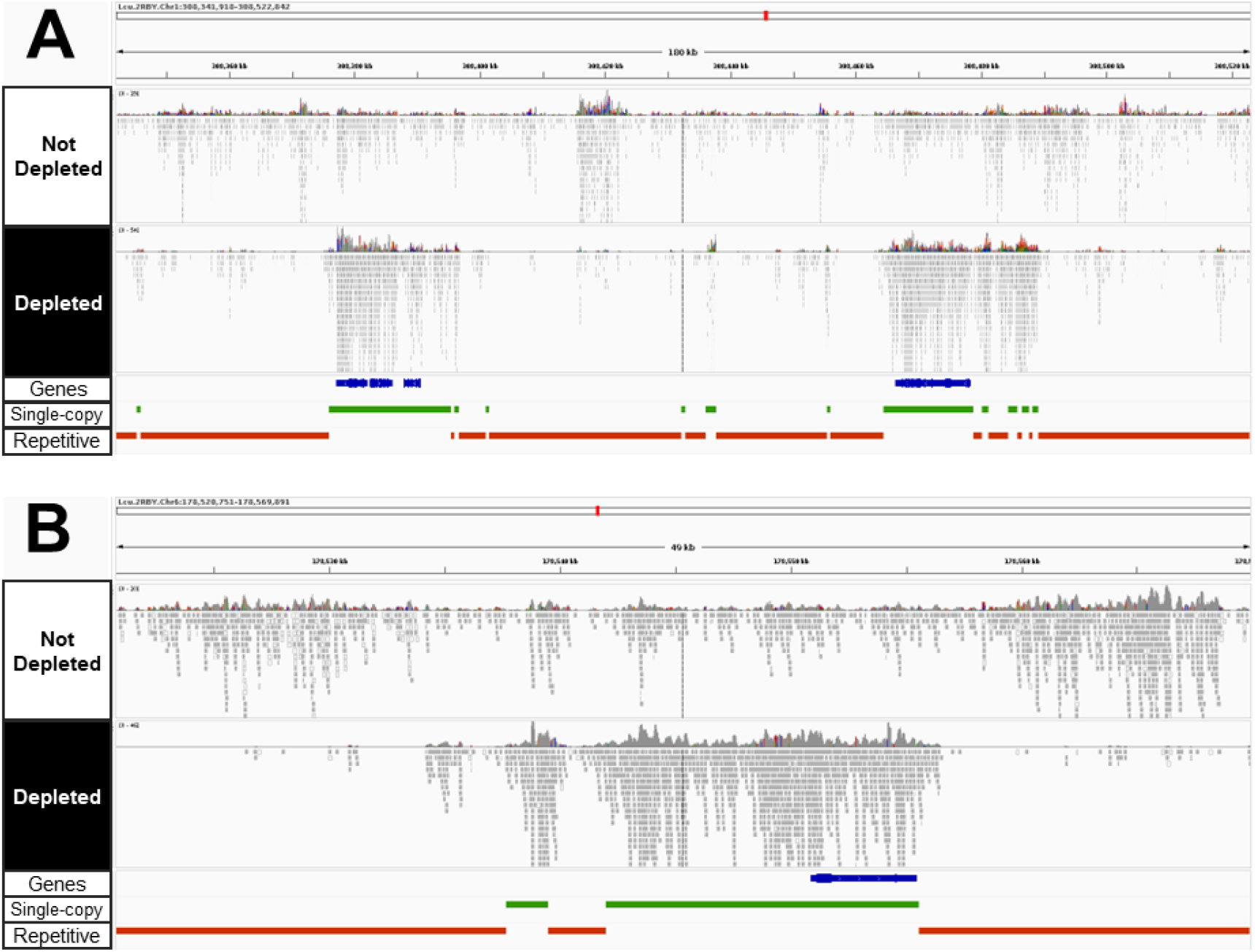
Reads mapped to repetitive or single-copy regions of the lentil genome with or without CRISPR/Cas9-mediated repeat depletion. Integrative Genome Browser Visualization (IGV) of Illumina sequencing data mapped to two representative genomic sites of ∼180 and ∼50 kbp. Tracks in blue, green and red represent annotated genes, single-copy regions and repetitive regions, respectively.

### Efficiency of CRISPR/Cas9-mediated depletion for different classes of nuclear repeats

We next examined the depletion of different classes of repetitive sequences in the *L. culinaris* genome. Reads mapping to the most abundant retroelements, namely the Ty3-Gypsy family (64% of the genome (Ramsay et al. 2021), were reduced by 47% in the depleted libraries, whereas those mapping to the Ty3-Copia family (15% of the genome) and other long terminal repeat (LTR) elements (3% of the genome) were depleted by 3% and 38%, respectively (**Figure 3A** and **Supplemental Table S5**). In contrast, there was no decrease in the abundance of other transposable elements (Line, CACTA, mu, hAT, Helitron, Harbinger, mariner and Sine), each representing < 1% of the genome (**Supplemental Table S5**), and there was no reduction in the number of reads mapping to tandem repeats (0.4% of the genome) or simple sequence repeats (8% of the genome) (**Figure 3A**). We observed a significant correlation between the variation in mapped reads after depletion and the abundance of these repeat classes in terms of overall repeat length and occurrence in the genome (**Figure 3B-C** and **Supplemental Table S5**). Given that the number of gRNAs targeting each repeat class increased proportionally with the repeat size and occurrence, the most efficiently depleted repetitive elements also featured a higher density of gRNA targets (**Figure 3D**). There was a significant correlation between the variation of mapping reads following depletion and the gRNA density over the whole target region when considering each single cut site in the genome (**Supplemental Figure S1**). In particular, target regions with a density > 8 gRNAs/kbp showed a read reduction in 85% of cases (**Supplemental Figure S1**) whereas regions targeted by < 8 gRNAs/kbp usually showed limited or no depletion (**Supplemental Figure S1**). We therefore concluded that the depletion efficiency across different repeat classes was dependent on the density of gRNA targets.

**Figure 3.**
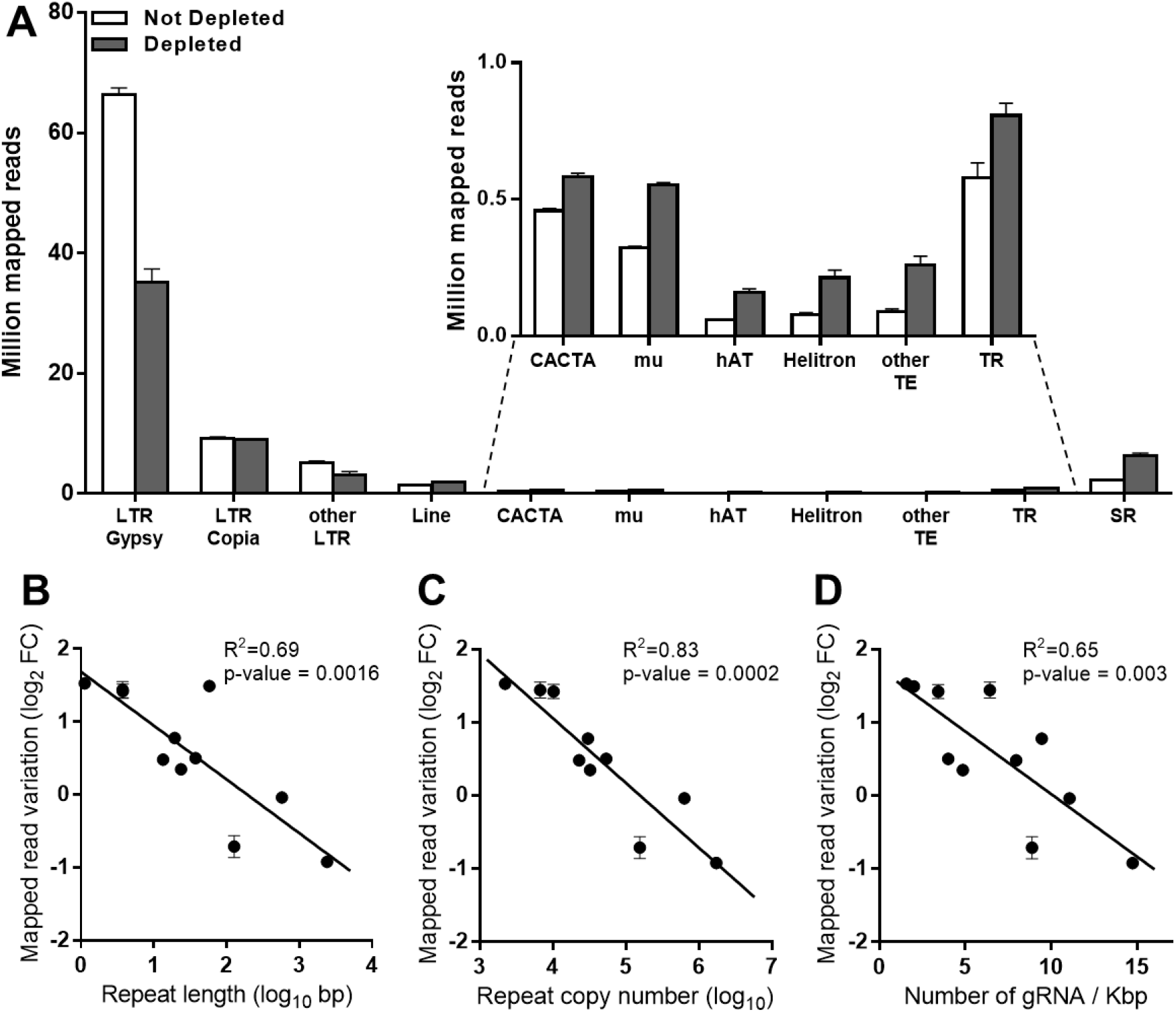
Sequencing data distribution after CRISPR/Cas9-mediated repeat depletion according to the class of nuclear repeat. (**A**) Number of reads (in millions) mapping to different nuclear repeat classes with or without CRISPR/Cas9-mediated repeat depletion: Ty3-Gypsy (LTR Gypsy), Ty1-Copia (LTR Copia), other LTR, Line, CACTA, mu, hAT, Helitron transposons, other transposable elements with abundance < 0.1% (Harbinger, mariner, Sine), tandem repeats (TR) and simple repeats (SR). Correlation between the variation of mapped reads following CRISPR/Cas9-mediated repeat depletion on the same repeat classes versus the repeat length (**B**), repeat copy number (**C**) and density of gRNAs targeting each repeat class (**D**). Data are means ± SE (n = 3). FC – fold change.

### Impact of CRISPR/Cas9-mediated repeat depletion on genotyping accuracy

Next, we investigated the impact of CRISPR/Cas9-mediated repeat depletion on the number of genomic positions in the single-copy regions where a base can be reliably genotyped (PASS at a depth of ≥ 5 reads). For this analysis, sequencing data generated from depleted and non-depleted *L. culinaris* cv. Castelluccio samples were downsampled from 62 to 6 million fragments to mimic lcWGS and ultra-lcWGS. Consistently more bases were genotyped within the single-copy regions of the depleted samples over the whole range considered (from ∼3.5 to ∼13-fold), with the highest gains at the lowest amounts of sequencing data (6 to 25 million fragments) (**Figure 4A**). Because a genotyped position does not necessarily allow the variant to be identified (this also depends on allele coverage), we also determined the impact of repeat depletion on variant calling. Following depletion, the total number of variants identified in the single-copy regions increased significantly from ∼4.5 to ∼18-fold, with a delta of ∼25,000 to ∼1 million more variants identified in the depleted sample (**Figure 4B**). Also the number of heterozygous variants identified was increased, although these constituted a minor fraction of total variants, as lentil is an autogamous species (**Figure 4C**). This allowed us to identify from ∼650 up to ∼50,000 variant positions that would be erroneously classified as reference without the depletion (**Figure 4D**). These false negative variants in the non-depleted sample were not called due to allelic imbalance caused by the low coverage (**Supplemental Figure S2**). Consistently, most of these false negative variants rescued by depletion were heterozygous (∼97%, **Figure 4D**).

**Figure 4.**
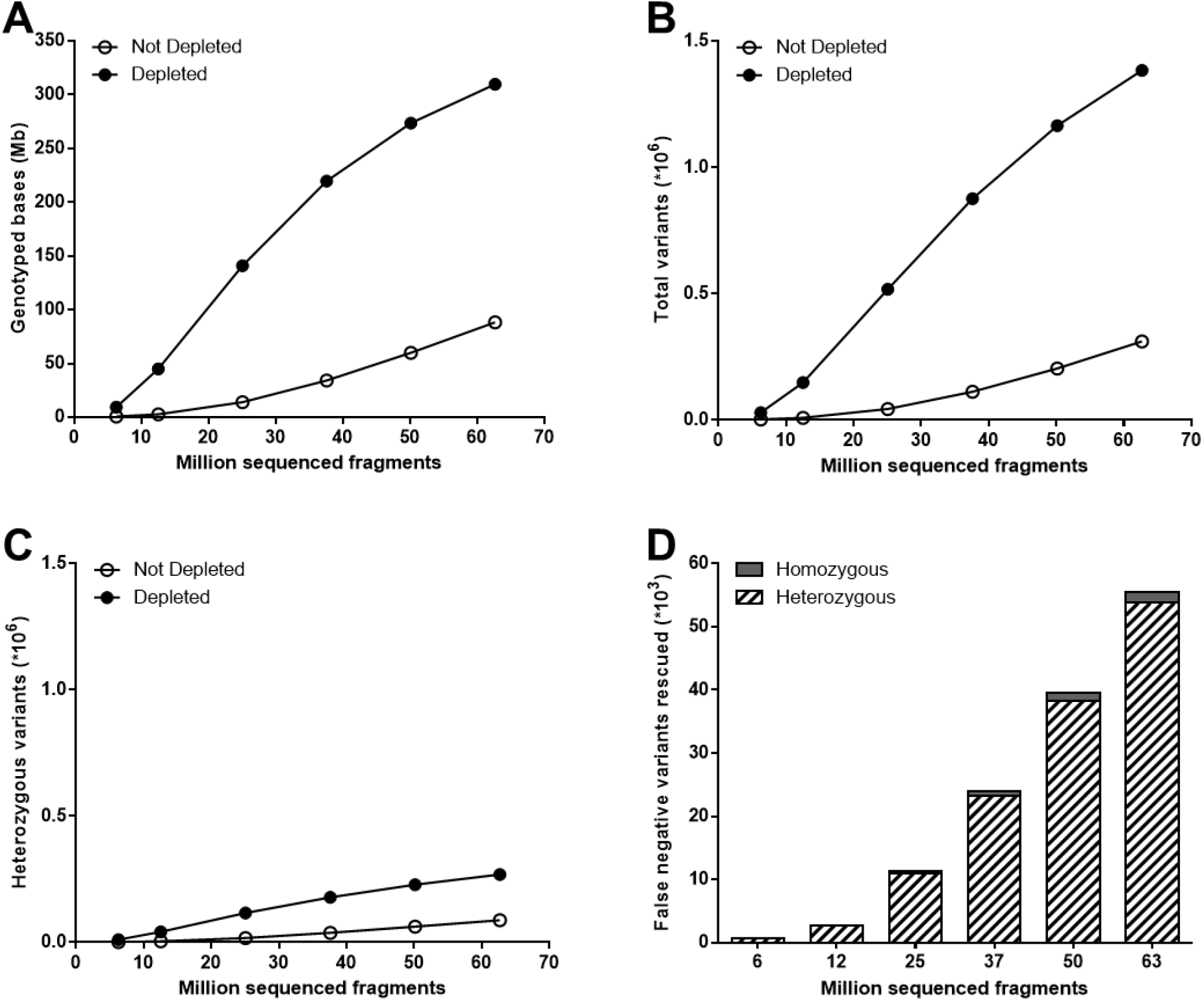
Genotyping performance on single-copy regions with or without CRISPR/Cas9-mediated repeat depletion starting from different amounts of sequencing data. (**A**) Number of genotyped positions. (**B**) Number of total variants identified. (**C**) Number of heterozygous variants identified. (**D**) Number of variants that were genotyped as reference (0/0) in the non-depleted sample and identified as homozygous (1/1) or heterozygous (1/0) alternative in the depleted sample. N=3 at 6, 12 and 25 million fragments, N=2 at 37, 50 and 63 million fragments.

### Performance of CRISPR/Cas9-mediated repeat depletion on different samples and library types

Finally, we assessed the performance of CRISPR/Cas9-mediated repeat depletion on different lentil genotypes, multiplexing levels and library types (**Supplemental Table S3** and **Supplemental Table S4**). Similar variations in the coverage of repetitive/single-copy regions and the number of mapped reads were observed when depleting multiplex libraries generated from a different cultivar (RB, Redberry) or from the closely-related species *L. orientalis* (**Figure 5A-B**) when compared to the original Castelluccio cultivar (**Figure 1**). To assess the impact of depletion when comparing different samples, these data were downsampled as described above, genotyped positions were identified and intersected with those of Castelluccio samples. Depletion improved the genotyping reproducibility, as the number of genotyped positions in common between all analyzed samples, within the single-copy regions, was consistently higher than in the condition without depletion (**Figure 5C**). Finally, there was no significant difference in the performance of CRISPR/Cas9-mediated repeat depletion when library multiplexing was increased from 8-plex to 96-plex while maintaining the 1:2.5 Cas9:gRNA ratio and 1 ng of treated library per sample (**Figure 6A-B**), or when treating standard singleplex WGS libraries (**Figure 6C-D**). Overall, these results demonstrated that CRISPR/Cas9-mediated repeat depletion using the same gRNA set is at least equally effective when applied to a group of two lentil cultivars and one close wild relative, providing an improved genotyping reproducibility and the possibility to process individual samples or multiple samples simultaneously.

**Figure 5.**
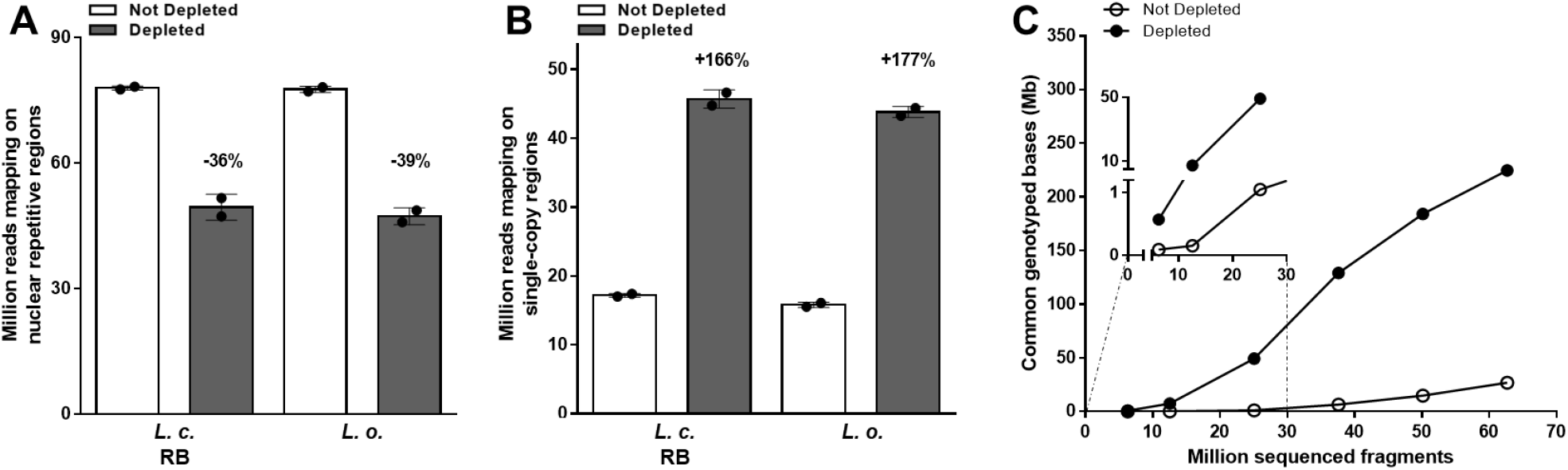
Performances of CRISPR/Cas9-mediated repeat depletion in different lentil samples. (**A-B**) Sequencing reads mapping to the repetitive or single-copy regions with or without CRISPR/Cas9-mediated repeat depletion in multiplex libraries generated from different lentil samples, namely *L. culinaris* cv. Redberry (*L. c*. RB) or *L. orientalis* (*L. o*.). Data are means ± SE (n = 2) normalized for the same sequencing input (50 million fragments). Variation percentages observed following CRISPR/Cas9-mediated repeat depletion are reported above each condition. (**C**) Number of genotyped positions in common between all samples analyzed in the study (3 distinct samples of *L. culinaris* cv. Castelluccio, 1 sample of *L. culinaris* cv. Redberry, and 1 samples of *L. orientalis*), within the single-copy regions. N=5 at 6, 12 and 25 million fragments, N=4 at 37, 50 and 63 million fragments.

**Figure 6.**
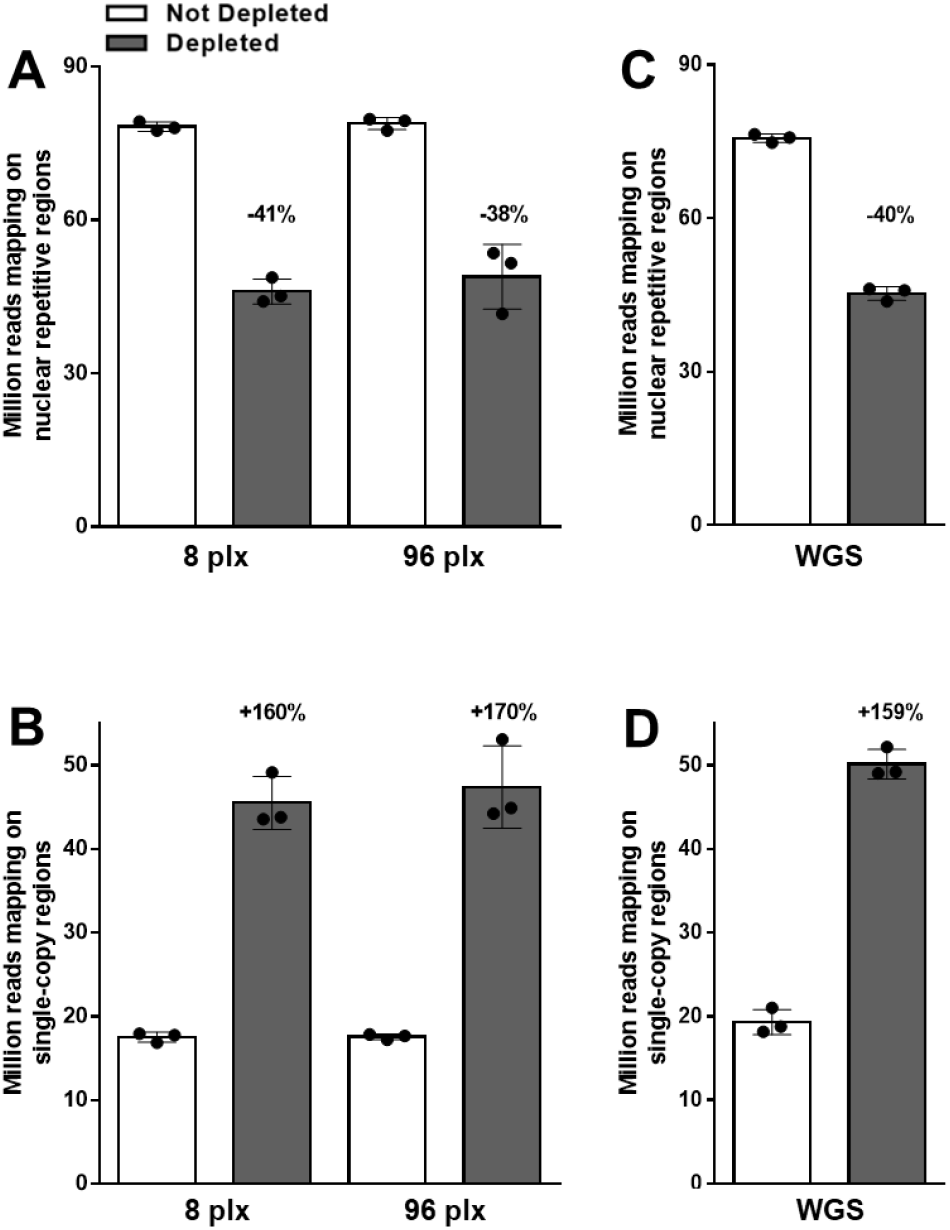
Performances of CRISPR/Cas9-mediated repeat depletion at different multiplexing level and library types. Sequencing reads mapping to the repetitive or single-copy regions with or without CRISPR/Cas9-mediated repeat depletion in (**A-B**) multiplex libraries containing 8 or 96 samples (plx) or (**E-F)** standard singleplex WGS libraries generated from one *L. culinaris* cv. Castelluccio, one *L. culinaris* cv. Redberry and one L. orientalis sample. Data are means ± SE (n = 3) normalized for the same sequencing input (50 million fragments). Variation percentages observed following CRISPR/Cas9mediated repeat depletion are reported above each condition.

## DISCUSSION

In a typical genotyping experiment based on sequencing, the data derived from repetitive DNA is directly proportional to the repeat content of the genome, however these data are largely uninformative. The bigger the genome, the more sequencing costs are therefore wasted on repeats. The traditional solution is the capture and sequencing of coding regions (WES). However, we approached the problem from the opposite perspective by depleting repetitive elements from the large genome of *L. culinaris* using CRISPR/Cas9 technology. Similar methods have been used for RNA-Seq library normalization (Prezza et al. 2020), metatrascriptomics (Gu et al. 2016), pathogen detection (Gu et al. 2016; Le et al. 2021) and single-cell analysis (Homberger et al. 2023; Le et al. 2021). Here, the main challenge is that 84% of the 3.7-Gb *L. culinaris* genome is repetitive DNA. The design of the gRNA array, therefore, required stringent multi-step filtering resulting in a set of ∼566,766 gRNAs. To our knowledge, this is the first time CRISPR/Cas9 has been used with such a large number of gRNAs either *in vitro* or *in vivo*, representing a fundamental advance in the technology platform beyond the specific goals of our project. Despite these technical challenges, CRISPR/Cas9-mediated repeat depletion produced sequencing libraries with consistently lower proportions of repetitive DNA (41.2% depletion) and enriched the singlecopy regions (160% increase), thus allowing the generation of more meaningful sequencing data.

Repetitive DNA in plant genomes can be divided into two broad categories: dispersed mobile elements and tandem/simple repeats. Dispersed mobile elements are made up of DNA transposons and retrotransposons, the most abundant of which are the LTR retrotransposons (Bennetzen and Wang 2014). Although mobile elements are not under the same selection pressure as genes, the degree of conservation across multiple copies of the same element is sufficiently high to allow the targeting of multiple copies with single gRNAs. The most efficient depletion (47%) was achieved for the most abundant LTR retrotransposon family (Gypsy, ∼64% of the genome), followed by all the other LTR families (–15%). The cutting of LTR elements by Cas9 was responsible for almost the entire depletion observed at the genome-wide level, whereas the depletion of other mobile elements was negligible. This was probably due to the lower repetition of such elements, which translated into a poor cutting frequency in the final gRNA design, comprising only gRNAs with 25 targets. These targets usually featured < 8 gRNA/kbp, namely a density associated with a low depletion rate. A similar observation was reported for RNA-Seq libraries, where a gRNA every 50–100 nucleotides (10–20 gRNAs/kbp) achieves excellent depletion results (Gu et al. 2016). Simple and tandem repeats were also not depleted efficiently, although the gRNA density was close to 8. In these cases, the highly repetitive motifs of such sequences may have reduced the cutting efficiency of Cas9 (Müller Paul et al. 2022). Given that LTR retrotransposons make up the majority of repeats in plant genomes (94% in *L. culinaris*) and are the principal cause of plant genome size variation (Bennetzen and Wang 2014; Lee and Kim 2014), the design of gRNAs to target only LTR sequences may be the most efficient strategy to reduce the genome size in sequencing experiments. Future gRNA designs in other species and further optimization of the *L. culinaris* design should maximize the gRNA number on these most abundant elements, instead of dispersing the effort across the remaining repetitive fraction (< 10%). Another factor influencing the efficiency of CRISPR/Cas9-mediated repeat depletion was the dose of Cas9 and gRNAs; doubling their dose indeed improved repeat depletion by ∼10%, albeit with a slight increase in overall costs. Finally, also the order of gRNA addition was important, as higher depletion efficiency was achieved when splitting gRNAs into three groups based on cutting frequency and using the groups sequentially. A possible explanation of this phenomenon is that most abundant gRNAs, with the highest number of target sites in the genome, could interfere with the “genome patrolling” of other RNPs targeting less abundant elements. Therefore, the gRNA target density, RNP concentration and prioritization of gRNAs with less abundant targets are factors that can improve depletion efficiency.

Organelle genomes often constitute a large fraction of DNA derived from plants (Sakamoto and Takami 2018). Although the mitochondrial and chloroplast genomes are smaller than the nuclear genome, they are present in multiple copies per cell, and they can represent > 20% of the total sequence data (Gargiulo et al. 2021; Ren et al. 2021). CRISPR/Cas9-mediated repeat depletion has been shown to reduce the fraction of sequencing libraries derived from organelle genomes in ATAC-Seq experiments (Montefiori et al. 2017). Our method was efficient for the depletion of chloroplast DNA (by 67%) while the depletion of mitochondrial DNA was only marginal, possibly reflecting the different abundance of the two organelle genomes in the starting genomic sample. Still, in lentil, the fraction of sequencing data attributable to organelles was largely due to chloroplasts (88%), whose depletion was therefore sufficient to decrease the data mapping on organelles after Cas9 treatment (−65% overall). Although in the case of lentil the total sequencing data attributable to organelles was rather low (∼1.3% in the not depleted libraries), the depletion of organelle DNA from sequencing libraries will be highly beneficial for organisms with a strongly unbalanced ratio of organelle vs nuclear DNA, such as *Cypripedium calceolus* (Gargiulo et al. 2021) and *Haematococcus pluvialis* (Ren et al. 2021). Although this has not been investigated in the present work, it is plausible that gRNAs designed for organelle’s genomes could also target nuclear integrants of plastid/mitochondrial DNA (NUPTs and NUMTs), that are homologous to the cpDNA and mtDNA (Zhang et al. 2020; Sloan et al. 2018).Although the efficiency of CRISPR/Cas9-mediated repeat depletion could be improved, the current set of gRNAs allowed us to genotype consistently more bases on single-copy as compared to not depleted libraries, and consequently to identify more genetic variants. We observed that coupling the depletion with 25 million sequencing fragments provided the best balance between costs and fraction of genotyped bases. In this condition, depleted samples reached an average coverage of ∼5 fold on single-copy regions, with gains of 10-and 12-fold in the number of genotyped positions and identified variants, respectively, as compared to not depleted libraries. We estimated that one would need 3.5-4 times more sequencing data to achieve the same performances when using not depleted libraries. By targeting the majority of genomic length (84%), the depletion allowed to pour a large fraction of sequencing data on the single-copy regions, that are instead just a small fraction (16%). For this reason, the CRISPR/Cas9-repeat depletion was more effective to improve genotyping performances than increasing the overall sequencing coverage, being also beneficial for the identification of heterozygous variants that would otherwise be missed due to unbalanced allelic sampling. Although this involved only ∼3% of total variants identified in lentil, as this is an autogamous diploid species, allelic imbalance is a well-known cause of errors in genotyping experiments based on sequencing (Cooke et al. 2016). CRISPR/Cas9-mediated repeat depletion can therefore improve the accuracy of genotyping experiments, especially in plants with highly heterozygous genomes and/or in polyploids.

Given that CRISPR/Cas9 repeat depletion allows to concentrate the sequencing data on the desired regions, the amount of positions that were genotyped in common between multiple samples was consistently higher in the depleted ones. In population studies, the shift in the distribution of sequencing data may contribute to reduce the number of missing data, thereby detecting a larger number of differences between samples. Furthermore, this approach was successful in different cultivars of *L. culinaris*, and also in the closely-related species *L. orientalis*. This is important because genotyping experiments typically include distant/wild relatives and related species, from which it is possible to develop evolutionary studies and plan breeding experiments, including the introgression of characters of interest. For example, the INCREASE project features a collection of 2000 lentil accessions that includes both cultivated varieties and local landraces (Guerra-García et al. 2021). Further experiments could determine whether the gRNAs designed in this study are also suitable for the depletion of repeats in other closely related leguminous species (Fabaceae) with very large and repetitive genomes, such as pea (*P. sativum*, 3.92 Gb, 83% repetitive)(Kreplak et al. 2019) and faba bean (*Vicia faba* L., 12 Gb, 79% transposon-derived repeats)(Jayakodi et al. 2023).

The cost of NGS library preparation for a genotyping project can easily exceed the cost of sequencing in the case of small genomes and/or ultra-lcWGS, especially given the steadily falling price of sequencing. More recent library-preparation kits circumvent several lengthy steps that require expensive reagents, and allow large sample sets to be processed in multiplex reactions. We used the Twist 96-Plex Library Prep Kit (formerly iGenomX Riptide) that constructs Illumina NGS libraries by polymerase-mediated extension of barcoded random primers. This type of library is beneficial for genotyping in general because random priming reduces uniform genome coverage but allows more reproducible sampling of the same sites across multiple samples (Siddique et al. 2019). Most importantly, the kit is designed to process large numbers of samples (up to 96 simultaneously) at low costs and without advanced equipment (just a multichannel pipette). To design, filter and synthesize the gRNA set targeting *L. culinaris* repeats required 10 weeks and cost ∼20,000 USD. The latter comprised gRNAs and depletion reagents sufficient for 30 reactions, each for a maximum of 96 samples treated in multiplex, corresponding to approximately 10 USD per sample. We estimated that the net cost to achieve ∼5-fold average coverage of the single-copy regions of the lentil genome by combining CRISPR/Cas9-mediated repeat depletion with a 96-plex Twist library is approximately US$75, made up of US$15 for library preparation, US$10 for depletion and US$40 for sequencing on a NovaSeq6000 S4 flowcell, generating 25 million fragments per sample. As such, our results demonstrated that depleted libraries are more informative than standard ones when normalized for the amount of sequencing data. Alternatively, CRISPR/Cas9-mediated repeat depletion can be used to reduce sequencing costs (by ∼75% in lentil), because the same number of genotyped bases (or detected variants) in single-copy regions can be detected with much less sequencing data. CRISPR/Cas9-mediated repeat depletion combined with Twist multiplex libraries is therefore an effective strategy for genotyping projects involving hundreds or thousands of samples. Dealing with less-repetitive datasets can also reduce the complexity of the genotyping analysis and the computational resources required.

The method therefore has the potential to increase our genetic knowledge of plant species that are currently difficult to analyze without a significant economic investment due to the large genome size and high proportion of repetitive DNA. Population studies, eQTL analysis, GWAS and pre-breeding programs are just some of the approaches that can benefit from CRISPR/Cas9-mediated repeat depletion.

## METHODS

### Multiplex-library preparation and sequencing

We prepared 8-plex and 96-plex multiplex libraries according to the Twist 96-Plex Library Preparation Kit protocol (Twist Bioscience, South San Francisco, CA, USA) with the following modifications. For each sample, we denatured 100 ng of genomic DNA (25 ng/µl) at 98 °C for 1 min. Ultra-low (30%) GC random primer set A was used for the extension and termination reaction (Reaction A) followed by 8 and 9 cycles of PCR amplification for the 96-plex and 8-plex libraries, respectively. Final libraries were purified using Twist DNA Purification Beads (0.65x volume) and a second round of purification was applied to the supernatant using 10 µl of beads to achieve a median insert size of 500 bp. Libraries were quantified using the Qubit BR DNA kit and a Qubit device (Thermo Fisher Scientific, Waltham, MA, USA) and size distributions were assessed using a Tape Station System (Agilent Technologies, Santa Clara, CA, USA). Non-depleted libraries were pooled at equimolar concentrations and sequenced on a NovaSeq6000 instrument (Illumina, San Diego, CA, USA) to generate 150-bp paired-end reads.

### WGS library preparation and sequencing

Genomic DNA samples were fragmented using a Covaris sonicator to achieve an average size of 400 bp, and Illumina PCR-free libraries were prepared from 700 ng DNA using the KAPA Hyper prep kit and unique dual-indexed adapters (5 µL of a 15 µM stock) according to the supplier’s protocol (Roche, Basel, Switzerland). The library concentration and size distribution were assessed on a Bioanalyzer (Agilent Technologies, Santa Clara, CA, USA). Non-depleted WGS libraries were pooled at equimolar concentrations and sequenced on a NovaSeq6000 instrument (Illumina, San Diego, CA, USA) to generate 150-bp paired-end reads.

### Design of gRNAs

The gRNA set was designed by JumpCode Genomics (San Diego, CA, USA) against the repetitive regions of the *L. culinaris* CDC Redberry v2.0 reference genome (Ramsay et al. 2021) (https://knowpulse.usask.ca/genome-assembly/Lcu.2RBY). The available repeat annotation (transposable elements and tandem repeats) was integrated with the annotation of simple and tandem repeats identified by RepeatMasker v4.0.6 and Tandem Repeat Finder v4.9 using 2 7 7 80 10 50 2000 -d -h parameters to identify intervals for gRNA design (**Supplemental Table S1**). Adjacent or overlapping intervals were collapsed into single intervals before design. As a first step, all 20 nt sequences with adjacent PAM sites for Cas9 (NGG) were identified in the target intervals. Second, the guides were filtered to exclude secondary structure, high and low GC content, homopolymers, dinucleotide repeats and low *in vitro* cleavage efficiency prediction scores (Azimuth algorithm; (Doench et al. 2016)). Third, the resulting guides were filtered to minimize off-target cleavage in single-copy regions of the genome by excluding guides that have complementary sites in genomic regions corresponding to genes and open-chromatin regions identified by ATAC-Seq (PRJNA912311) (allowing for up to 3 mismatches). As a final step, and to reduce the number of guides in the set, guides were selected to have no fewer than 25 cleavage sites each and to maintain an inter-guide spacing of at least 500 bp. The final guide set, comprising 569,088 unique guides, was split into 11 pools for the purpose of synthesis. The number of copies of each guide varied and reflected the number of on-target cleavage sites for each guide. DNA oligonucleotides containing the target-specific 20 nt gRNA sequence and invariant single gRNA sequence were synthesized, after which pools of oligonucleotides were amplified by PCR and converted to RNA by *in vitro* transcription. The products of transcription were treated with DNase I and column purified to generate the final gRNA material. Pools 1–3, 5–8 and 10 contain only gRNAs targeting the nuclear genome. Pools 9 and 11 contain both nuclear and chloroplast genome gRNAs, and pool 4 is the only pool containing gRNAs that target the nuclear, chloroplast and mitochondrial genomes (**Supplemental Table S2** and **Supplemental File S1**). The number of gRNAs targeting each repeat class are reported in **Supplemental Table S5**.

### Repeat depletion with JumpCode CRISPRclean

Repetitive regions were depleted using the Cas9 protein and the custom gRNA set described above, according to the Jumpcode CRISPRclean Ribosomal RNA Depletion from Human RNA-Seq Libraries for Illumina Sequencing protocol (Jumpcode Genomics, San Diego, CA, USA) with the following modifications. The input was 10 and 100 ng for the 8-plex and 96-plex libraries, respectively. Depletion was carried out either using all gRNA simultaneously or by splitting the gRNA pools into three groups based on cutting frequency, which were used sequentially in order of increasing cutting frequency (**Supplemental Table S2**). The sequential depletion strategy was also conducted using the double amounts of gRNAs and Cas9. The reaction volume was 20 µl when using all gRNAs simultaneously or 26 µl for the sequential and double sequential protocols. The quantity of each gRNA pool per reaction is shown in **Supplemental Table S2** and amounted to 620 ng in the simultaneous and sequential depletion reactions or 1240 ng in the double sequential depletion reaction. The Cas9 enzyme was diluted 1:5 in 1x Cas9 Buffer and 0.0029 µl was used per ng gRNA. The reactions were incubated at 37 °C and libraries were treated in the presence of gRNAs for a total of 1 h (simultaneous depletion protocol) or 3 h (sequential and double sequential depletion protocols, with the sequential gRNA pools added at 1-h intervals). The depleted samples were then size selected using 0.6x volume of AMPure XP Beads (Beckman Coulter, Brea, CA, USA). Libraries were amplified with 10 and 6 PCR cycles for the 8-plex and 96-plex libraries, respectively before final purification with 60µl (0.6x volume) AMPure XP Beads. The concentrations of depleted libraries were measured using the Qubit system and size distributions were assessed on a Tape Station System as described above. Depleted libraries were pooled at equimolar concentrations and sequenced on a NovaSeq6000 instrument to generate 150-bp paired-end reads.

### Data analysis and variant calling

Raw read quality was assessed using FastQC (http://www.bioinformatics.babraham.ac.uk/projects/fastqc/) and the multiplex libraries were demultiplexed using fgbio v1.3.0 DemuxFastqs, assigning fragments by exploiting the unique sample identifier included during first-strand synthesis. The raw reads were then quality filtered and the Illumina sequencing adapters removed using scythe v0.991 and syckle v1.33, respectively. Filtered reads were aligned to the *L. culinaris* v2.0 reference genome using bwa-mem v2.2.1 and the resulting alignments were converted to bam files and sorted using samtools v1.13. PCR-derived duplicates were removed using the GATK MarkDuplicates tool v4.1.7.0 and overlapping portions of the paired-end reads were clipped using the fgbio v1.3.0 ClipBam tool. The resulting bam files were used to calculate coverage depth, breadth and fraction of PASS bases (at ≥ 5x) using bedtools v2.30.0 genomecov and GATK v3.8 CallableLoci, respectively. The number of reads aligning to the reference genome and to different regions of interest was calculated using samtools v1.13 with option -c to discard reads with a 2308 sam flag in order to consider only the primary alignment, thus omitting repetitive counts of the same multimapping reads. When necessary, sequencing data were normalized to a pre-defined number of input fragments using seqtk sample v1.3.

The variation of mapped reads in depleted *vs* not depleted libraries was calculated using the formula:

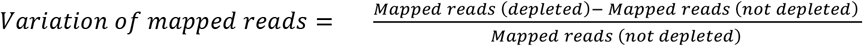

To achieve a normal distribution of variation, in Supplemental Figure S1 the variation of mapped read coverage between the depleted and not depleted libraries was calculated using the following formula:

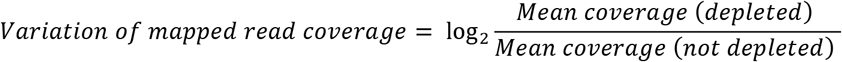

Regions with zero coverage in either depleted or not depleted conditions (6,097 and 6,319, respectively, over a total of 237,136 repetitive segments, of which 4,662 in common) were excluded from the calculation, as they represented a negligible fraction of total (3.3%) and showed a minimal coverage also in the opposite condition (< 0.2 fold). The plot in Supplemental Figure S1 was generated using the ggplot package in R.

Genomic variants were identified using GATK HaplotypeCaller v4.1.7.0 with the parameters “--min-base-quality-score 20 -ERC GVCF”. Individual gVCF files were merged using GATK GenomicsDBImport v4.1.7.0 and the final VCF file was generated using GATK GenotypeGVCFs v4.1.7.0. Variant filtration was achieved using GATK hard filters (https://gatk.broadinstitute.org/hc/en-us/articles/360037499012?id=3225).

## DATA ACCESS

The sequencing datasets generated and analysed in this study have been submitted to the NCBI GenBank repository (https://www.ncbi.nlm.nih.gov/bioproject/) under accession number PRJNA915594 and PRJNA912311.

## ACKNOWLEDGMENTS

This research was supported by the European Union’s Horizon 2020 research and innovation program, through the project INCREASE (www.pulsesincrease.eu) (grant agreement No. 862862). The founding sponsors had no role in the design of the study; in the collection, analyses, or interpretation of data; in the writing of the manuscript; and in the decision to publish the results. We acknowledge Matteo De Biasi for the support in bioinformatic analysis, the Twist Bioscience and Jumpcode Genomics teams for the excellent technical support.

## Authors’ contributions

Conceptualization MR, MD and RP; Methodology MR, LM and MD; Software LM and GL; Investigation LDA, EBE, GC and FL; Formal analysis MR, LM and LDA; Validation GF, LN and LV; Resources KB, LR and DJK; Data Curation LM and GL; Writing – Original Draft MR; Writing – Review & Editing LM, LDA and MD; Visualization MR and LM; Supervision MR and MD; Project administration MR; Funding acquisition EBI, MD and RP.

