## Supplemental Tables for "CRISPR/Cas9-based repeat depletion for the high-throughput genotyping of complex plant genomes"

**Supplemental Table S1. Genome assembly, annotation and gRNA design statistics**

The genome assembly and annotation statistics are presented for the *L. culinaris* reference genome Lcu.2RBY (CDC Redberry) v2.0. The available annotation was integrated with the simple and tandem repeat identification (see Methods). The reported features of the gRNAs include the regions used as input for the design: the on-target regions (repeats), corresponding to the total repeats (transposable elements, tandem and simple repeats) after collapsing adjacent or overlapping intervals into single intervals, the off-target regions (single-copy regions) comprising genes, putative regulatory regions identified based on ATAC-Seq data, and all the remaining genomic regions not included in the repetitive regions. The final gRNA targets in the nuclear, mitochondrial and chloroplast genomes were calculated by combining all gRNA binding sites plus 500 bp upstream and downstream, representing the size of the genomic libraries in this project. Both for genomic DNA and gRNA design, the fraction of each feature is calculated over the total size of the nuclear or organelle genome. N/A, not applicable.

| **GENOMIC FEATURES** | **bp** | **genomic fraction** |
|---|---:|---:|
| Genome | 3,760,304,314 | N/A |
| Transposable Elements (TE) | 3,103,065,790 | 82.5% |
| Simple Repeats (SR) | 58,089,181 | 1.5% |
| Tandem Repeats (TR) | 13,460,124 | 0.4% |
| Total repeats (merging TE+SR+TR) | 3,162,675,774 | 84.1% |
| Genes | 185,392,774 | 4.9% |
| Putative Regulatory Elements (ATAC-seq) | 78,309,986 | 2.1% |
| Mitochondrial genome | 489,326 | N/A |
| Chloroplast genome | 118,064 | N/A |

| **gRNA DESIGN FEATURES** | **bp** | **genomic fraction** |
|---|---:|---:|
| Off-target regions (single-copy regions) | 563,967,150 | 15% |
| Target regions (repetitive regions) | 3,195,729,774 | 85% |
| Final gRNA target design on nuclear genome | 2,904,915,709 | 77% |
| Final gRNA target design on Mitochondrial genome | 422,811 | 86% |
| Final gRNA target design on Chloroplast genome | 118,064 | 100% |

**Supplemental Table S2. Composition of gRNA pools.**

For each pool, the table shows the number of total gRNAs, the genomic target, the relative frequency, quantity (ng) per reaction, and the subdivision of pools into three groups based on cutting frequency. The relative frequency is the ratio between the number of cut sites for each gRNA and the minimum number of cut sites (25). N/A, not applicable, as the exact number of organelle genomes is not known.

| Pool | Genomic target | Total gRNAs | Relative frequency | ng per reaction (1x) | Depletion group (based on cutting frequency) |
|---|---|---|---|---|---|
| 1 | nuclear | 60,000 | 1 | 40 | Group 1 |
| 2 | nuclear | 60,000 | 1 | 40 | Group 1 |
| 3 | nuclear | 60,000 | 1 | 40 | Group 1 |
| 4 | nuclear | 39,463 | 1 | 60 | Group 2 |
| | mitochondrial | 7,480 | N/A | | |
| | chloroplast | 13,050 | N/A | | |
| 5 | nuclear | 60,000 | 2 | 40 | Group 1 |
| 6 | nuclear | 60,000 | 2 | 40 | Group 2 |
| 7 | nuclear | 49,574 | 2 | 40 | Group 2 |
| | nuclear | 7,394 | 3 | | |
| 8 | nuclear | 60,000 | 3 | 40 | Group 2 |
| 9 | nuclear | 34,198 | 4 | 80 | Group 3 |
| | nuclear | 20,486 | 5 | | |
| | chloroplast | 5,000 | N/A | | |
| 10 | nuclear | 33,613 | 6-9 | 120 | Group 3 |
| | nuclear | 25,695 | 20-99 | | |
| 11 | nuclear | 50,397 | 10-19 | 80 | Group 3 |
| | nuclear | 2,700 | ≥100 | | |
| | chloroplast | 6,850 | N/A | | |

**Supplemental Table S3. Composition of 8-plex and 96-plex multiplex libraries**

For each multiplex library, the table reports the ID of the library and its samples, indicating the species and the cultivar.
Each single replicate is associated with its internal barcode.
castel, cv. Castelluccio; RB, cv. Redberry; rep, replicate

| **Twist 8 plx libraries from *L. culinaris* cv. Castelluccio** | | | | | | **depletion condition** |
|---|---|---|---|---|---|---|
| **1092** | | **1146** | | **4589** | | |
| **sample ID** | **barcode** | **sample ID** | **barcode** | **sample ID** | **barcode** | |
| L_culinaris_castel_3650_rep1 | GCTTAACG | L_culinaris_castel_3650_rep1 | CGTACGTA | L_culinaris_castel_3650_rep1 | AGCTAGCT | Twist 8 plx - L. culinaris cv. Castelluccio - Not Depleted |
| L_culinaris_castel_3650_rep2 | GTACCAAC | L_culinaris_castel_3650_rep2 | AGGAACGT | L_culinaris_castel_3650_rep2 | AGGTAGGT | |
| L_culinaris_castel_3650_rep3 | GAAGTCCT | L_culinaris_castel_3650_rep3 | CTAGCTAG | L_culinaris_castel_3650_rep3 | CAGTCAGT | |
| L_culinaris_castel_3652_rep1 | CCAATACG | L_culinaris_castel_3652_rep1 | CTCTCAGT | L_culinaris_castel_3652_rep1 | CAAGCTTG | |
| L_culinaris_castel_3652_rep2 | AGCTCTAG | L_culinaris_castel_3652_rep2 | GTACTGCA | L_culinaris_castel_3652_rep2 | AACCTTGG | |
| L_culinaris_castel_3652_rep3 | CGTAGCTA | L_culinaris_castel_3652_rep3 | CTTCCATG | L_culinaris_castel_3652_rep3 | ATCGTAGC | |
| L_culinaris_castel_316_rep1 | CTCTCACA | L_culinaris_castel_316_rep1 | TACGAACC | L_culinaris_castel_316_rep1 | CACTAGAC | |
| L_culinaris_castel_316_rep2 | TGACCACA | L_culinaris_castel_316_rep2 | TACCATGG | L_culinaris_castel_316_rep2 | GCTTCCTA | |
| **1145** | | **1091** | | **1306** | | |
| **sample ID** | **barcode** | **sample ID** | **barcode** | **sample ID** | **barcode** | |
| lent_castel_3650_1_rep1 | GCTTAACG | lent_castel_3650_1 | CGTACGTA | L_culinaris_castel_3650_rep1 | AGCTAGCT | Twist 8 plx - L. culinaris cv. Castelluccio - Depleted, All gRNA |
| lent_castel_3650_1_rep2 | GTACCAAC | lent_castel_3650_1 | AGGAACGT | L_culinaris_castel_3650_rep2 | AGGTAGGT | |
| lent_castel_3650_1_rep3 | GAAGTCCT | lent_castel_3650_1 | CTAGCTAG | L_culinaris_castel_3650_rep3 | CAGTCAGT | |
| lent_castel_3652_1_rep1 | CCAATACG | lent_castel_3652_1 | CTCTCAGT | L_culinaris_castel_3652_rep1 | CAAGCTTG | |
| lent_castel_3652_1_rep2 | AGCTCTAG | lent_castel_3652_1 | GTACTGCA | L_culinaris_castel_3652_rep2 | AACCTTGG | |
| lent_castel_3652_1_rep3 | CGTAGCTA | lent_castel_3652_1 | CTTCCATG | L_culinaris_castel_3652_rep3 | ATCGTAGC | |
| L_culinaris_castel_316_rep1 | CTCTCACA | L_culinaris_castel_316_rep1 | TACGAACC | L_culinaris_castel_316_rep1 | CACTAGAC | |
| L_culinaris_castel_316_rep2 | TGACCACA | L_culinaris_castel_316_rep2 | TACCATGG | L_culinaris_castel_316_rep2 | GCTTCCTA | |
| **2971** | | **1144** | | **5700** | | |
| **sample ID** | **barcode** | **sample ID** | **barcode** | **sample ID** | **barcode** | |
| L_culinaris_castel_3650_rep1 | GCTTAACG | lent_castel_3650_1_rep1 | CGTACGTA | lent_castel_3650_rep1 | AGCTAGCT | Twist 8 plx - L. culinaris cv. Castelluccio - Depleted, 3grp gRNA |
| L_culinaris_castel_3650_rep2 | GTACCAAC | lent_castel_3650_1_rep2 | AGGAACGT | lent_castel_3650_rep2 | AGGTAGGT | |
| L_culinaris_castel_3650_rep3 | GAAGTCCT | lent_castel_3650_1_rep3 | CTAGCTAG | lent_castel_3650_rep3 | CAGTCAGT | |
| L_culinaris_castel_3652_rep1 | CCAATACG | lent_castel_3652_1_rep1 | CTCTCAGT | lent_castel_3652_1_rep1 | CAAGCTTG | |
| L_culinaris_castel_3652_rep2 | AGCTCTAG | lent_castel_3652_1_rep2 | GTACTGCA | lent_castel_3652_1_rep2 | AACCTTGG | |
| L_culinaris_castel_3652_rep3 | CGTAGCTA | lent_castel_3652_1_rep3 | CTTCCATG | lent_castel_3652_1_rep3 | ATCGTAGC | |
| L_culinaris_castel_316_rep1 | CTCTCACA | L_culinaris_castel_316_rep1 | TACGAACC | L_culinaris_castel_316_rep1 | CACTAGAC | |
| L_culinaris_castel_316_rep2 | TGACCACA | L_culinaris_castel_316_rep2 | TACCATGG | L_culinaris_castel_316_rep2 | GCTTCCTA | |
| **4017** | | **1308** | | **5701** | | |
| **sample ID** | **barcode** | **sample ID** | **barcode** | **sample ID** | **barcode** | |
| lent_castel_3650_1_rep1 | GCTTAACG | lent_castel_3650_1_rep1 | CGTACGTA | lent_castel_3650_rep1 | AGCTAGCT | Twist 8 plx - L. culinaris cv. Castelluccio - Depleted, 3grp 2x gRNA |
| lent_castel_3650_1_rep2 | GTACCAAC | lent_castel_3650_1_rep2 | AGGAACGT | lent_castel_3650_rep2 | AGGTAGGT | |
| lent_castel_3650_1_rep3 | GAAGTCCT | lent_castel_3650_1_rep3 | CTAGCTAG | lent_castel_3650_rep3 | CAGTCAGT | |
| lent_castel_3652_1_rep4 | CCAATACG | lent_castel_3652_1_rep1 | CTCTCAGT | lent_castel_3652_1_rep1 | CAAGCTTG | |
| lent_castel_3652_1_rep5 | AGCTCTAG | lent_castel_3652_1_rep2 | GTACTGCA | lent_castel_3652_1_rep2 | AACCTTGG | |
| lent_castel_3652_1_rep6 | CGTAGCTA | lent_castel_3652_1_rep3 | CTTCCATG | lent_castel_3652_1_rep3 | ATCGTAGC | |
| L_culinaris_castel_316_rep1 | CTCTCACA | L_culinaris_castel_316_rep1 | TACGAACC | L_culinaris_castel_316_rep1 | CACTAGAC | |
| L_culinaris_castel_316_rep2 | TGACCACA | L_culinaris_castel_316_rep2 | TACCATGG | L_culinaris_castel_316_rep2 | GCTTCCTA | |

| **Twist 8 plx libraries from *L. culinaris* cv. Redberry and *L. orientalis*** | | | | | | | | **depletion condition** |
|---|---|---|---|---|---|---|---|---|
| **1288** | | **1290** | | | | | | |
| **sample ID** | **barcode** | **sample ID** | **barcode** | | | | | |
| Sallys_Redberry_lens_culinaris_rep1 | ATCCGGTA | Sallys_Redberry_lens_culinaris_rep1 | GCTACGAT | | | | | Twist 8 plx - L. culinaris cv. Redberry and L. orientalis - Not Depleted |
| Sallys_Redberry_lens_culinaris_rep2 | GACATCAC | Sallys_Redberry_lens_culinaris_rep2 | CTAGAGCT | | | | | |
| Sallys_Redberry_lens_culinaris_rep3 | TCGATGGT | Sallys_Redberry_lens_culinaris_rep3 | ACTGACTG | | | | | |
| Sallys_Redberry_lens_culinaris_rep4 | CTGTCAGA | Sallys_Redberry_lens_culinaris_rep4 | GGAAGCAT | | | | | |
| IG_72529_Lens_Orientalis_rep1 | TGGTACGT | IG_72529_Lens_Orientalis_rep1 | CCATATGG | | | | | |
| IG_72529_Lens_Orientalis_rep2 | GATGCATC | IG_72529_Lens_Orientalis_rep2 | CAGTTGAC | | | | | |
| IG_72529_Lens_Orientalis_rep3 | CTTCCAAC | IG_72529_Lens_Orientalis_rep3 | ACACGTCA | | | | | |
| IG_72529_Lens_Orientalis_rep4 | ACACGTGT | IG_72529_Lens_Orientalis_rep4 | ACCAACGT | | | | | |
| **4884** | | **5685** | | **5684** | | **2970** | | |
| **sample ID** | **barcode** | **sample ID** | **barcode** | **sample ID** | **barcode** | **sample ID** | **barcode** | |
| Sally_s_Redberry_lens_culinaris_rep1 | ATCCGGTA | Sally_s_Redberry_lens_culinaris_rep1 | ATCCGGTA | Sally_s_Redberry_lens_culinaris_rep1 | ATCCGGTA | Sally_s_Redberry_lens_cu | ATCCGGTA | |

| | | | | | | | | |
|---|---|---|---|---|---|---|---|---|
| Sally_s_Redberry_lens_culinaris_rep2 | GACATCAC | Sally_s_Redberry_lens_culinaris_rep2 | GACATCAC | Sally_s_Redberry_lens_culinaris_rep2 | GACATCAC | Sally_s_Redberry_lens_cu | GACATCAC | |
| Sally_s_Redberry_lens_culinaris_rep3 | TCGATGGT | Sally_s_Redberry_lens_culinaris_rep3 | TCGATGGT | Sally_s_Redberry_lens_culinaris_rep3 | TCGATGGT | Sally_s_Redberry_lens_cu | TCGATGGT | Twist 8 plx - L. culinaris cv. Redberry and L. orientalis - Depleted, 3grp 2x gRNA |
| Sally_s_Redberry_lens_culinaris_rep4 | CTGTCAGA | Sally_s_Redberry_lens_culinaris_rep4 | CTGTCAGA | Sally_s_Redberry_lens_culinaris_rep4 | CTGTCAGA | Sally_s_Redberry_lens_cu | CTGTCAGA | |
| IG_72529_Lens_Orientalis_rep1 | TGGTACGT | IG_72529_Lens_Orientalis_rep1 | TGGTACGT | IG_72529_Lens_Orientalis_rep1 | TGGTACGT | IG_72529_Lens_Orientalis | TGGTACGT | |
| IG_72529_Lens_Orientalis_rep2 | GATGCATC | IG_72529_Lens_Orientalis_rep2 | GATGCATC | IG_72529_Lens_Orientalis_rep2 | GATGCATC | IG_72529_Lens_Orientalis | GATGCATC | |
| IG_72529_Lens_Orientalis_rep3 | CTTCCAAC | IG_72529_Lens_Orientalis_rep3 | CTTCCAAC | IG_72529_Lens_Orientalis_rep3 | CTTCCAAC | IG_72529_Lens_Orientalis | CTTCCAAC | |
| IG_72529_Lens_Orientalis_rep4 | ACACGTGT | IG_72529_Lens_Orientalis_rep4 | ACACGTGT | IG_72529_Lens_Orientalis_rep4 | ACACGTGT | IG_72529_Lens_Orientalis | ACACGTGT | |

| **Twist 96 plx libraries from *L. culinaris* cv. Castelluccio and Redberry** | | | | | | **depletion condition** |
|---|---|---|---|---|---|---|
| **1090** | | **4560** | | **4588** | | **Not Depleted** |
| **sample ID** | **barcode** | **sample ID** | **barcode** | **sample ID** | **barcode** | |
| lent_castel_3648_1 | CGTACGTA | Sally_s_Redberry_lens_culinaris_rep1 | CGTACGTA | lent_castel_3648_1 | CGTACGTA | |
| lent_castel_3648_1 | AGCTAGCT | Sally_s_Redberry_lens_culinaris_rep2 | AGGAACGT | lent_castel_3648_1 | AGCTAGCT | |
| lent_castel_3648_1 | GCTTAACG | Sally_s_Redberry_lens_culinaris_rep3 | CTAGCTAG | lent_castel_3648_1 | GCTTAACG | |
| lent_castel_3648_1 | ATCCGGTA | Sally_s_Redberry_lens_culinaris_rep4 | CTCTCAGT | lent_castel_3648_1 | ATCCGGTA | |
| lent_castel_3648_1 | AGGAACGT | Sally_s_Redberry_lens_culinaris_rep5 | GTACTGCA | lent_castel_3648_1 | AGGAACGT | |
| lent_castel_3648_1 | AGGTAGGT | Sally_s_Redberry_lens_culinaris_rep6 | CTTCCATG | lent_castel_3648_1 | AGGTAGGT | |
| lent_castel_3648_1 | GTACCAAC | Sally_s_Redberry_lens_culinaris_rep7 | TACGAACC | lent_castel_3648_1 | GTACCAAC | |
| lent_castel_3648_1 | GACATCAC | Sally_s_Redberry_lens_culinaris_rep8 | TACCATGG | lent_castel_3648_1 | GACATCAC | |
| lent_castel_3648_1 | CTAGCTAG | Sally_s_Redberry_lens_culinaris_rep9 | AGCTAGCT | lent_castel_3648_1 | CTAGCTAG | |
| lent_castel_3648_1 | CAGTCAGT | Sally_s_Redberry_lens_culinaris_rep10 | AGGTAGGT | lent_castel_3648_1 | CAGTCAGT | |
| lent_castel_3648_1 | GAAGTCCT | Sally_s_Redberry_lens_culinaris_rep11 | CAGTCAGT | lent_castel_3648_1 | GAAGTCCT | |
| lent_castel_3648_1 | TCGATGGT | Sally_s_Redberry_lens_culinaris_rep12 | CAAGCTTG | lent_castel_3648_1 | TCGATGGT | |
| lent_castel_3648_1 | CTCTCAGT | Sally_s_Redberry_lens_culinaris_rep13 | AACCTTGG | lent_castel_3648_1 | CTCTCAGT | |
| lent_castel_3648_1 | CAAGCTTG | Sally_s_Redberry_lens_culinaris_rep14 | ATCGTAGC | lent_castel_3648_1 | CAAGCTTG | |
| lent_castel_3648_1 | CCAATACG | Sally_s_Redberry_lens_culinaris_rep15 | CACTAGAC | lent_castel_3648_1 | CCAATACG | |
| lent_castel_3648_1 | CTGTCAGA | Sally_s_Redberry_lens_culinaris_rep16 | GCTTCCTA | lent_castel_3648_1 | CTGTCAGA | |
| lent_castel_3648_1 | GTACTGCA | Sally_s_Redberry_lens_culinaris_rep17 | GCTTAACG | lent_castel_3648_1 | GTACTGCA | |
| lent_castel_3648_1 | AACCTTGG | Sally_s_Redberry_lens_culinaris_rep18 | GTACCAAC | lent_castel_3648_1 | AACCTTGG | |
| lent_castel_3648_1 | AGCTCTAG | Sally_s_Redberry_lens_culinaris_rep19 | GAAGTCCT | lent_castel_3648_1 | AGCTCTAG | |
| lent_castel_3648_1 | TGGTACGT | Sally_s_Redberry_lens_culinaris_rep20 | CCAATACG | lent_castel_3648_1 | TGGTACGT | |
| lent_castel_3648_1 | CTTCCATG | Sally_s_Redberry_lens_culinaris_rep21 | AGCTCTAG | lent_castel_3648_1 | CTTCCATG | |
| lent_castel_3648_1 | ATCGTAGC | Sally_s_Redberry_lens_culinaris_rep22 | CGTAGCTA | lent_castel_3648_1 | ATCGTAGC | |
| lent_castel_3648_1 | CGTAGCTA | Sally_s_Redberry_lens_culinaris_rep23 | CTCTCACA | lent_castel_3648_1 | CGTAGCTA | |
| lent_castel_3648_1 | GATGCATC | Sally_s_Redberry_lens_culinaris_rep24 | TGACCACA | lent_castel_3648_1 | GATGCATC | |
| lent_castel_3648_1 | TACGAACC | Sally_s_Redberry_lens_culinaris_rep25 | ATCCGGTA | lent_castel_3648_1 | TACGAACC | |
| lent_castel_3648_1 | CACTAGAC | Sally_s_Redberry_lens_culinaris_rep26 | GACATCAC | lent_castel_3648_1 | CACTAGAC | |
| lent_castel_3648_1 | CTCTCACA | Sally_s_Redberry_lens_culinaris_rep27 | TCGATGGT | lent_castel_3648_1 | CTCTCACA | |
| lent_castel_3648_1 | CTTCCAAC | Sally_s_Redberry_lens_culinaris_rep28 | CTGTCAGA | lent_castel_3648_1 | CTTCCAAC | |
| lent_castel_3648_1 | TACCATGG | Sally_s_Redberry_lens_culinaris_rep29 | TGGTACGT | lent_castel_3648_1 | TACCATGG | |
| lent_castel_3648_1 | GCTTCCTA | Sally_s_Redberry_lens_culinaris_rep30 | GATGCATC | lent_castel_3648_1 | GCTTCCTA | |
| lent_castel_3648_1 | TGACCACA | Sally_s_Redberry_lens_culinaris_rep31 | CTTCCAAC | lent_castel_3648_1 | TGACCACA | |
| lent_castel_3648_1 | ACACGTGT | Sally_s_Redberry_lens_culinaris_rep32 | ACACGTGT | lent_castel_3648_1 | ACACGTGT | |
| lent_castel_3649_1 | TAGGCCAT | Sally_s_Redberry_lens_culinaris_rep33 | TAGGCCAT | lent_castel_3649_1 | TAGGCCAT | |
| lent_castel_3649_1 | GCTACGAT | Sally_s_Redberry_lens_culinaris_rep34 | CTGTACAG | lent_castel_3649_1 | GCTACGAT | |
| lent_castel_3649_1 | GATCAGCT | Sally_s_Redberry_lens_culinaris_rep35 | TCGAGTTG | lent_castel_3649_1 | GATCAGCT | |
| lent_castel_3649_1 | AACGTTGC | Sally_s_Redberry_lens_culinaris_rep36 | AAGGTTCC | lent_castel_3649_1 | AACGTTGC | |
| lent_castel_3649_1 | CTGTACAG | Sally_s_Redberry_lens_culinaris_rep37 | GCCGTATA | lent_castel_3649_1 | CTGTACAG | |
| lent_castel_3649_1 | CTAGAGCT | Sally_s_Redberry_lens_culinaris_rep38 | ACCTGTTC | lent_castel_3649_1 | CTAGAGCT | |
| lent_castel_3649_1 | CGTAGCAT | Sally_s_Redberry_lens_culinaris_rep39 | ACAGTCAC | lent_castel_3649_1 | CGTAGCAT | |
| lent_castel_3649_1 | TGCACAAC | Sally_s_Redberry_lens_culinaris_rep40 | GAGTTCTG | lent_castel_3649_1 | TGCACAAC | |
| lent_castel_3649_1 | TCGAGTTG | Sally_s_Redberry_lens_culinaris_rep41 | GCTACGAT | lent_castel_3649_1 | TCGAGTTG | |
| lent_castel_3649_1 | ACTGACTG | Sally_s_Redberry_lens_culinaris_rep42 | CTAGAGCT | lent_castel_3649_1 | ACTGACTG | |
| lent_castel_3649_1 | GTGTTGAC | Sally_s_Redberry_lens_culinaris_rep43 | ACTGACTG | lent_castel_3649_1 | GTGTTGAC | |
| lent_castel_3649_1 | CTCTACAC | Sally_s_Redberry_lens_culinaris_rep44 | GGAAGCAT | lent_castel_3649_1 | CTCTACAC | |
| lent_castel_3649_1 | AAGGTTCC | Sally_s_Redberry_lens_culinaris_rep45 | CCATATGG | lent_castel_3649_1 | AAGGTTCC | |
| lent_castel_3649_1 | GGAAGCAT | Sally_s_Redberry_lens_culinaris_rep46 | CAGTTGAC | lent_castel_3649_1 | GGAAGCAT | |
| lent_castel_3649_1 | AGTGTCTG | Sally_s_Redberry_lens_culinaris_rep47 | ACACGTCA | lent_castel_3649_1 | AGTGTCTG | |
| lent_castel_3649_1 | GCTATTCC | Sally_s_Redberry_lens_culinaris_rep48 | ACCAACGT | lent_castel_3649_1 | GCTATTCC | |
| lent_castel_3649_1 | GCCGTATA | L_culinaris_castel_316 | GATCAGCT | lent_castel_3649_1 | GCCGTATA | |
| lent_castel_3649_1 | CCATATGG | L_culinaris_castel_316 | CGTAGCAT | lent_castel_3649_1 | CCATATGG | |

| | | | | | | |
|---|---|---|---|---|---|---|
| lent_castel_3649_1 | TGGTCATG | L_culinaris_castel_316 | GTGTTGAC | lent_castel_3649_1 | TGGTCATG | |
| lent_castel_3649_1 | CCTAATCC | L_culinaris_castel_316 | AGTGTCTG | lent_castel_3649_1 | CCTAATCC | |
| lent_castel_3649_1 | ACCTGTTC | L_culinaris_castel_316 | TGGTCATG | lent_castel_3649_1 | ACCTGTTC | |
| lent_castel_3649_1 | CAGTTGAC | L_culinaris_castel_316 | ACTGTGAC | lent_castel_3649_1 | CAGTTGAC | |
| lent_castel_3649_1 | ACTGTGAC | L_culinaris_castel_316 | CTTGTCCA | lent_castel_3649_1 | ACTGTGAC | |
| lent_castel_3649_1 | CATGCTTC | L_culinaris_castel_316 | CGATCGAT | lent_castel_3649_1 | CATGCTTC | |
| lent_castel_3649_1 | ACAGTCAC | L_culinaris_castel_316 | AACGTTGC | lent_castel_3649_1 | ACAGTCAC | |
| lent_castel_3649_1 | ACACGTCA | L_culinaris_castel_316 | TGCACAAC | lent_castel_3649_1 | ACACGTCA | |
| lent_castel_3649_1 | CTTGTCCA | L_culinaris_castel_316 | CTCTACAC | lent_castel_3649_1 | CTTGTCCA | |
| lent_castel_3649_1 | TGACGTGT | L_culinaris_castel_316 | GCTATTCC | lent_castel_3649_1 | TGACGTGT | |
| lent_castel_3649_1 | GAGTTCTG | L_culinaris_castel_316 | CCTAATCC | lent_castel_3649_1 | GAGTTCTG | |
| lent_castel_3649_1 | ACCAACGT | L_culinaris_castel_316 | CATGCTTC | lent_castel_3649_1 | ACCAACGT | |
| lent_castel_3649_1 | CGATCGAT | L_culinaris_castel_316 | TGACGTGT | lent_castel_3649_1 | CGATCGAT | |
| lent_castel_3649_1 | ACCACATG | L_culinaris_castel_316 | ACCACATG | lent_castel_3649_1 | ACCACATG | |
| L_culinaris_castel_316 | GACTTCAG | L_culinaris_castel_316 | GACTTCAG | L_culinaris_castel_316 | GACTTCAG | |
| L_culinaris_castel_316 | GAGTTCAC | L_culinaris_castel_316 | ACCAAGGA | L_culinaris_castel_316 | GAGTTCAC | |
| L_culinaris_castel_316 | GGATTAGG | L_culinaris_castel_316 | GCCGTTAA | L_culinaris_castel_316 | GGATTAGG | |
| L_culinaris_castel_316 | TGGTCTAG | L_culinaris_castel_316 | TGACTGAC | L_culinaris_castel_316 | TGGTCTAG | |
| L_culinaris_castel_316 | ACCAAGGA | L_culinaris_castel_316 | TGGTGTAC | L_culinaris_castel_316 | ACCAAGGA | |
| L_culinaris_castel_316 | GCGCTATA | L_culinaris_castel_316 | GTACAGCT | L_culinaris_castel_316 | GCGCTATA | |
| L_culinaris_castel_316 | GAACCATC | L_culinaris_castel_316 | GGTTCCAA | L_culinaris_castel_316 | GAACCATC | |
| L_culinaris_castel_316 | CTACCATC | L_culinaris_castel_316 | GAGACACA | L_culinaris_castel_316 | CTACCATC | |
| L_culinaris_castel_316 | GCCGTTAA | L_culinaris_castel_316 | GAGTTCAC | L_culinaris_castel_316 | GCCGTTAA | |
| L_culinaris_castel_316 | GTGAACAG | L_culinaris_castel_316 | GCGCTATA | L_culinaris_castel_316 | GTGAACAG | |
| L_culinaris_castel_316 | ATGCATGC | L_culinaris_castel_316 | GTGAACAG | L_culinaris_castel_316 | ATGCATGC | |
| L_culinaris_castel_316 | TCCAAGGT | L_culinaris_castel_316 | TGCACTAG | L_culinaris_castel_316 | TCCAAGGT | |
| L_culinaris_castel_316 | TGACTGAC | L_culinaris_castel_316 | AACCGGTT | L_culinaris_castel_316 | TGACTGAC | |
| L_culinaris_castel_316 | TGCACTAG | L_culinaris_castel_316 | ATCCTAGG | L_culinaris_castel_316 | TGCACTAG | |
| L_culinaris_castel_316 | GATCCTAG | L_culinaris_castel_316 | CAACCTAG | L_culinaris_castel_316 | GATCCTAG | |
| L_culinaris_castel_316 | CTCATGAG | L_culinaris_castel_316 | AGTGGTCT | L_culinaris_castel_316 | CTCATGAG | |
| L_culinaris_castel_316 | TGGTGTAC | L_culinaris_castel_316 | GGATTAGG | L_culinaris_castel_316 | TGGTGTAC | |
| L_culinaris_castel_316 | AACCGGTT | L_culinaris_castel_316 | GAACCATC | L_culinaris_castel_316 | AACCGGTT | |
| L_culinaris_castel_316 | CAACGTAC | L_culinaris_castel_316 | ATGCATGC | L_culinaris_castel_316 | CAACGTAC | |
| L_culinaris_castel_316 | GAGACAGT | L_culinaris_castel_316 | GATCCTAG | L_culinaris_castel_316 | GAGACAGT | |
| L_culinaris_castel_316 | GTACAGCT | L_culinaris_castel_316 | CAACGTAC | L_culinaris_castel_316 | GTACAGCT | |
| L_culinaris_castel_316 | ATCCTAGG | L_culinaris_castel_316 | GAAGCTTC | L_culinaris_castel_316 | ATCCTAGG | |
| L_culinaris_castel_316 | GAAGCTTC | L_culinaris_castel_316 | TAGCTACG | L_culinaris_castel_316 | GAAGCTTC | |
| L_culinaris_castel_316 | GTGTCACA | L_culinaris_castel_316 | GAACTGCT | L_culinaris_castel_316 | GTGTCACA | |
| L_culinaris_castel_316 | GGTTCCAA | L_culinaris_castel_316 | TGGTCTAG | L_culinaris_castel_316 | GGTTCCAA | |
| L_culinaris_castel_316 | CAACCTAG | L_culinaris_castel_316 | CTACCATC | L_culinaris_castel_316 | CAACCTAG | |
| L_culinaris_castel_316 | TAGCTACG | L_culinaris_castel_316 | TCCAAGGT | L_culinaris_castel_316 | TAGCTACG | |
| L_culinaris_castel_316 | GACATCTG | L_culinaris_castel_316 | CTCATGAG | L_culinaris_castel_316 | GACATCTG | |
| L_culinaris_castel_316 | GAGACACA | L_culinaris_castel_316 | GAGACAGT | L_culinaris_castel_316 | GAGACACA | |
| L_culinaris_castel_316 | AGTGGTCT | L_culinaris_castel_316 | GTGTCACA | L_culinaris_castel_316 | AGTGGTCT | |
| L_culinaris_castel_316 | GAACTGCT | L_culinaris_castel_316 | GACATCTG | L_culinaris_castel_316 | GAACTGCT | |
| L_culinaris_castel_316 | CACATGTG | L_culinaris_castel_316 | CACATGTG | L_culinaris_castel_316 | CACATGTG | |
| **4014** | | **4890** | | **4886** | | **Depleted 3grp 2x gRNA** |
| **sample ID** | **barcode** | **sample ID** | **barcode** | **sample ID** | **barcode** | |
| lent_castel_3648_1_rep1 | CGTACGTA | Sally_s_Redberry_lens_culinaris_rep1 | CGTACGTA | lent_castel_3648_1_rep1 | CGTACGTA | |
| lent_castel_3648_1_rep10 | CAGTCAGT | Sally_s_Redberry_lens_culinaris_rep2 | AGGAACGT | lent_castel_3648_1_rep2 | AGCTAGCT | |
| lent_castel_3648_1_rep11 | GAAGTCCT | Sally_s_Redberry_lens_culinaris_rep3 | CTAGCTAG | lent_castel_3648_1_rep3 | GCTTAACG | |
| lent_castel_3648_1_rep12 | TCGATGGT | Sally_s_Redberry_lens_culinaris_rep4 | CTCTCAGT | lent_castel_3648_1_rep4 | ATCCGGTA | |
| lent_castel_3648_1_rep13 | CTCTCAGT | Sally_s_Redberry_lens_culinaris_rep5 | GTACTGCA | lent_castel_3649_1_rep1 | TAGGCCAT | |
| lent_castel_3648_1_rep14 | CAAGCTTG | Sally_s_Redberry_lens_culinaris_rep6 | CTTCCATG | lent_castel_3649_1_rep2 | GCTACGAT | |
| lent_castel_3648_1_rep15 | CCAATACG | Sally_s_Redberry_lens_culinaris_rep7 | TACGAACC | lent_castel_3649_1_rep3 | GATCAGCT | |
| lent_castel_3648_1_rep16 | CTGTCAGA | Sally_s_Redberry_lens_culinaris_rep8 | TACCATGG | lent_castel_3649_1_rep4 | AACGTTGC | |
| lent_castel_3648_1_rep17 | GTACTGCA | Sally_s_Redberry_lens_culinaris_rep9 | AGCTAGCT | L_culinaris_castel_316 | GACTTCAG | |
| lent_castel_3648_1_rep18 | AACCTTGG | Sally_s_Redberry_lens_culinaris_rep10 | AGGTAGGT | L_culinaris_castel_316 | GAGTTCAC | |
| lent_castel_3648_1_rep19 | AGCTCTAG | Sally_s_Redberry_lens_culinaris_rep11 | CAGTCAGT | L_culinaris_castel_316 | GGATTAGG | |
| lent_castel_3648_1_rep2 | AGCTAGCT | Sally_s_Redberry_lens_culinaris_rep12 | CAAGCTTG | L_culinaris_castel_316 | TGGTCTAG | |
| lent_castel_3648_1_rep20 | TGGTACGT | Sally_s_Redberry_lens_culinaris_rep13 | AACCTTGG | lent_castel_3648_1_rep5 | AGGAACGT | |

| | | | | | |
|---|---|---|---|---|---|
| lent_castel_3648_1_rep21 | CTTCCATG | Sally_s_Redberry_lens_culinaris_rep14 | ATCGTAGC | lent_castel_3648_1_rep6 | AGGTAGGT |
| lent_castel_3648_1_rep22 | ATCGTAGC | Sally_s_Redberry_lens_culinaris_rep15 | CACTAGAC | lent_castel_3648_1_rep7 | GTACCAAC |
| lent_castel_3648_1_rep23 | CGTAGCTA | Sally_s_Redberry_lens_culinaris_rep16 | GCTTCCTA | lent_castel_3648_1_rep8 | GACATCAC |
| lent_castel_3648_1_rep24 | GATGCATC | Sally_s_Redberry_lens_culinaris_rep17 | GCTTAACG | lent_castel_3649_1_rep5 | CTGTACAG |
| lent_castel_3648_1_rep25 | TACGAACC | Sally_s_Redberry_lens_culinaris_rep18 | GTACCAAC | lent_castel_3649_1_rep6 | CTAGAGCT |
| lent_castel_3648_1_rep26 | CACTAGAC | Sally_s_Redberry_lens_culinaris_rep19 | GAAGTCCT | lent_castel_3649_1_rep7 | CGTAGCAT |
| lent_castel_3648_1_rep27 | CTCTCACA | Sally_s_Redberry_lens_culinaris_rep20 | CCAATACG | lent_castel_3649_1_rep8 | TGCACAAC |
| lent_castel_3648_1_rep28 | CTTCCAAC | Sally_s_Redberry_lens_culinaris_rep21 | AGCTCTAG | L_culinaris_castel_316 | ACCAAGGA |
| lent_castel_3648_1_rep29 | TACCATGG | Sally_s_Redberry_lens_culinaris_rep22 | CGTAGCTA | L_culinaris_castel_316 | GCGCTATA |
| lent_castel_3648_1_rep3 | GCTTAACG | Sally_s_Redberry_lens_culinaris_rep23 | CTCTCACA | L_culinaris_castel_316 | GAACCATC |
| lent_castel_3648_1_rep30 | GCTTCCTA | Sally_s_Redberry_lens_culinaris_rep24 | TGACCACA | L_culinaris_castel_316 | CTACCATC |
| lent_castel_3648_1_rep31 | TGACCACA | Sally_s_Redberry_lens_culinaris_rep25 | ATCCGGTA | lent_castel_3648_1_rep9 | CTAGCTAG |
| lent_castel_3648_1_rep32 | ACACGTGT | Sally_s_Redberry_lens_culinaris_rep26 | GACATCAC | lent_castel_3648_1_rep10 | CAGTCAGT |
| lent_castel_3648_1_rep4 | ATCCGGTA | Sally_s_Redberry_lens_culinaris_rep27 | TCGATGGT | lent_castel_3648_1_rep11 | GAAGTCCT |
| lent_castel_3648_1_rep5 | AGGAACGT | Sally_s_Redberry_lens_culinaris_rep28 | CTGTCAGA | lent_castel_3648_1_rep12 | TCGATGGT |
| lent_castel_3648_1_rep6 | AGGTAGGT | Sally_s_Redberry_lens_culinaris_rep29 | TGGTACGT | lent_castel_3649_1_rep9 | TCGAGTTG |
| lent_castel_3648_1_rep7 | GTACCAAC | Sally_s_Redberry_lens_culinaris_rep30 | GATGCATC | lent_castel_3649_1_rep10 | ACTGACTG |
| lent_castel_3648_1_rep8 | GACATCAC | Sally_s_Redberry_lens_culinaris_rep31 | CTTCCAAC | lent_castel_3649_1_rep11 | GTGTTGAC |
| lent_castel_3648_1_rep9 | CTAGCTAG | Sally_s_Redberry_lens_culinaris_rep32 | ACACGTGT | lent_castel_3649_1_rep12 | CTCTACAC |
| lent_castel_3649_1_rep_17 | GCCGTATA | Sally_s_Redberry_lens_culinaris_rep33 | TAGGCCAT | L_culinaris_castel_316 | GCCGTTAA |
| lent_castel_3649_1_rep_18 | CCATATGG | Sally_s_Redberry_lens_culinaris_rep34 | CTGTACAG | L_culinaris_castel_316 | GTGAACAG |
| lent_castel_3649_1_rep_19 | TGGTCATG | Sally_s_Redberry_lens_culinaris_rep35 | TCGAGTTG | L_culinaris_castel_316 | ATGCATGC |
| lent_castel_3649_1_rep_20 | CCTAATCC | Sally_s_Redberry_lens_culinaris_rep36 | AAGGTTCC | L_culinaris_castel_316 | TCCAAGGT |
| lent_castel_3649_1_rep1 | TAGGCCAT | Sally_s_Redberry_lens_culinaris_rep37 | GCCGTATA | lent_castel_3648_1_rep13 | CTCTCAGT |
| lent_castel_3649_1_rep10 | ACTGACTG | Sally_s_Redberry_lens_culinaris_rep38 | ACCTGTTC | lent_castel_3648_1_rep14 | CAAGCTTG |
| lent_castel_3649_1_rep11 | GTGTTGAC | Sally_s_Redberry_lens_culinaris_rep39 | ACAGTCAC | lent_castel_3648_1_rep15 | CCAATACG |
| lent_castel_3649_1_rep12 | CTCTACAC | Sally_s_Redberry_lens_culinaris_rep40 | GAGTTCTG | lent_castel_3648_1_rep16 | CTGTCAGA |
| lent_castel_3649_1_rep13 | AAGGTTCC | Sally_s_Redberry_lens_culinaris_rep41 | GCTACGAT | lent_castel_3649_1_rep13 | AAGGTTCC |
| lent_castel_3649_1_rep14 | GGAAGCAT | Sally_s_Redberry_lens_culinaris_rep42 | CTAGAGCT | lent_castel_3649_1_rep14 | GGAAGCAT |
| lent_castel_3649_1_rep15 | AGTGTCTG | Sally_s_Redberry_lens_culinaris_rep43 | ACTGACTG | lent_castel_3649_1_rep15 | AGTGTCTG |
| lent_castel_3649_1_rep16 | GCTATTCC | Sally_s_Redberry_lens_culinaris_rep44 | GGAAGCAT | lent_castel_3649_1_rep16 | GCTATTCC |
| lent_castel_3649_1_rep2 | GCTACGAT | Sally_s_Redberry_lens_culinaris_rep45 | CCATATGG | L_culinaris_castel_316 | TGACTGAC |
| lent_castel_3649_1_rep21 | ACCTGTTC | Sally_s_Redberry_lens_culinaris_rep46 | CAGTTGAC | L_culinaris_castel_316 | TGCACTAG |
| lent_castel_3649_1_rep22 | CAGTTGAC | Sally_s_Redberry_lens_culinaris_rep47 | ACACGTCA | L_culinaris_castel_316 | GATCCTAG |
| lent_castel_3649_1_rep23 | ACTGTGAC | Sally_s_Redberry_lens_culinaris_rep48 | ACCAACGT | L_culinaris_castel_316 | CTCATGAG |
| lent_castel_3649_1_rep24 | CATGCTTC | L_culinaris_castel_316 | GATCAGCT | lent_castel_3648_1_rep17 | GTACTGCA |
| lent_castel_3649_1_rep25 | ACAGTCAC | L_culinaris_castel_316 | CGTAGCAT | lent_castel_3648_1_rep18 | AACCTTGG |
| lent_castel_3649_1_rep26 | ACACGTCA | L_culinaris_castel_316 | GTGTTGAC | lent_castel_3648_1_rep19 | AGCTCTAG |
| lent_castel_3649_1_rep27 | CTTGTCCA | L_culinaris_castel_316 | AGTGTCTG | lent_castel_3648_1_rep20 | TGGTACGT |
| lent_castel_3649_1_rep28 | TGACGTGT | L_culinaris_castel_316 | TGGTCATG | lent_castel_3649_1_rep_17 | GCCGTATA |
| lent_castel_3649_1_rep29 | GAGTTCTG | L_culinaris_castel_316 | ACTGTGAC | lent_castel_3649_1_rep_18 | CCATATGG |
| lent_castel_3649_1_rep3 | GATCAGCT | L_culinaris_castel_316 | CTTGTCCA | lent_castel_3649_1_rep_19 | TGGTCATG |
| lent_castel_3649_1_rep30 | ACCAACGT | L_culinaris_castel_316 | CGATCGAT | lent_castel_3649_1_rep_20 | CCTAATCC |
| lent_castel_3649_1_rep31 | CGATCGAT | L_culinaris_castel_316 | AACGTTGC | L_culinaris_castel_316 | TGGTGTAC |
| lent_castel_3649_1_rep32 | ACCACATG | L_culinaris_castel_316 | TGCACAAC | L_culinaris_castel_316 | AACCGGTT |
| lent_castel_3649_1_rep4 | AACGTTGC | L_culinaris_castel_316 | CTCTACAC | L_culinaris_castel_316 | CAACGTAC |
| lent_castel_3649_1_rep5 | CTGTACAG | L_culinaris_castel_316 | GCTATTCC | L_culinaris_castel_316 | GAGACAGT |
| lent_castel_3649_1_rep6 | CTAGAGCT | L_culinaris_castel_316 | CCTAATCC | lent_castel_3648_1_rep21 | CTTCCATG |
| lent_castel_3649_1_rep7 | CGTAGCAT | L_culinaris_castel_316 | CATGCTTC | lent_castel_3648_1_rep22 | ATCGTAGC |
| lent_castel_3649_1_rep8 | TGCACAAC | L_culinaris_castel_316 | TGACGTGT | lent_castel_3648_1_rep23 | CGTAGCTA |
| lent_castel_3649_1_rep9 | TCGAGTTG | L_culinaris_castel_316 | ACCACATG | lent_castel_3648_1_rep24 | GATGCATC |
| L_culinaris_castel_316 | GACTTCAG | L_culinaris_castel_316 | GACTTCAG | lent_castel_3649_1_rep21 | ACCTGTTC |
| L_culinaris_castel_316 | GTGAACAG | L_culinaris_castel_316 | ACCAAGGA | lent_castel_3649_1_rep22 | CAGTTGAC |
| L_culinaris_castel_316 | ATGCATGC | L_culinaris_castel_316 | GCCGTTAA | lent_castel_3649_1_rep23 | ACTGTGAC |
| L_culinaris_castel_316 | TCCAAGGT | L_culinaris_castel_316 | TGACTGAC | lent_castel_3649_1_rep24 | CATGCTTC |
| L_culinaris_castel_316 | TGACTGAC | L_culinaris_castel_316 | TGGTGTAC | L_culinaris_castel_316 | GTACAGCT |
| L_culinaris_castel_316 | TGCACTAG | L_culinaris_castel_316 | GTACAGCT | L_culinaris_castel_316 | ATCCTAGG |
| L_culinaris_castel_316 | GATCCTAG | L_culinaris_castel_316 | GGTTCCAA | L_culinaris_castel_316 | GAAGCTTC |
| L_culinaris_castel_316 | CTCATGAG | L_culinaris_castel_316 | GAGACACA | L_culinaris_castel_316 | GTGTCACA |
| L_culinaris_castel_316 | TGGTGTAC | L_culinaris_castel_316 | GAGTTCAC | lent_castel_3648_1_rep25 | TACGAACC |
| L_culinaris_castel_316 | AACCGGTT | L_culinaris_castel_316 | GCGCTATA | lent_castel_3648_1_rep26 | CACTAGAC |

| | | | | | |
|---|---|---|---|---|---|
| L_culinaris_castel_316 | CAACGTAC | L_culinaris_castel_316 | GTGAACAG | lent_castel_3648_1_rep27 | CTCTCACA |
| L_culinaris_castel_316 | GAGTTCAC | L_culinaris_castel_316 | TGCACTAG | lent_castel_3648_1_rep28 | CTTCCAAC |
| L_culinaris_castel_316 | GAGACAGT | L_culinaris_castel_316 | AACCGGTT | lent_castel_3649_1_rep25 | ACAGTCAC |
| L_culinaris_castel_316 | GTACAGCT | L_culinaris_castel_316 | ATCCTAGG | lent_castel_3649_1_rep26 | ACACGTCA |
| L_culinaris_castel_316 | ATCCTAGG | L_culinaris_castel_316 | CAACCTAG | lent_castel_3649_1_rep27 | CTTGTCCA |
| L_culinaris_castel_316 | GAAGCTTC | L_culinaris_castel_316 | AGTGGTCT | lent_castel_3649_1_rep28 | TGACGTGT |
| L_culinaris_castel_316 | GTGTCACA | L_culinaris_castel_316 | GGATTAGG | L_culinaris_castel_316 | GGTTCCAA |
| L_culinaris_castel_316 | GGTTCCAA | L_culinaris_castel_316 | GAACCATC | L_culinaris_castel_316 | CAACCTAG |
| L_culinaris_castel_316 | CAACCTAG | L_culinaris_castel_316 | ATGCATGC | L_culinaris_castel_316 | TAGCTACG |
| L_culinaris_castel_316 | TAGCTACG | L_culinaris_castel_316 | GATCCTAG | L_culinaris_castel_316 | GACATCTG |
| L_culinaris_castel_316 | GACATCTG | L_culinaris_castel_316 | CAACGTAC | lent_castel_3648_1_rep29 | TACCATGG |
| L_culinaris_castel_316 | GAGACACA | L_culinaris_castel_316 | GAAGCTTC | lent_castel_3648_1_rep30 | GCTTCCTA |
| L_culinaris_castel_316 | GGATTAGG | L_culinaris_castel_316 | TAGCTACG | lent_castel_3648_1_rep31 | TGACCACA |
| L_culinaris_castel_316 | AGTGGTCT | L_culinaris_castel_316 | GAACTGCT | lent_castel_3648_1_rep32 | ACACGTGT |
| L_culinaris_castel_316 | GAACTGCT | L_culinaris_castel_316 | TGGTCTAG | lent_castel_3649_1_rep29 | GAGTTCTG |
| L_culinaris_castel_316 | CACATGTG | L_culinaris_castel_316 | CTACCATC | lent_castel_3649_1_rep30 | ACCAACGT |
| L_culinaris_castel_316 | TGGTCTAG | L_culinaris_castel_316 | TCCAAGGT | lent_castel_3649_1_rep31 | CGATCGAT |
| L_culinaris_castel_316 | ACCAAGGA | L_culinaris_castel_316 | CTCATGAG | lent_castel_3649_1_rep32 | ACCACATG |
| L_culinaris_castel_316 | GCGCTATA | L_culinaris_castel_316 | GAGACAGT | L_culinaris_castel_316 | GAGACACA |
| L_culinaris_castel_316 | GAACCATC | L_culinaris_castel_316 | GTGTCACA | L_culinaris_castel_316 | AGTGGTCT |
| L_culinaris_castel_316 | CTACCATC | L_culinaris_castel_316 | GACATCTG | L_culinaris_castel_316 | GAACTGCT |
| L_culinaris_castel_316 | GCCGTTAA | L_culinaris_castel_316 | CACATGTG | L_culinaris_castel_316 | CACATGTG |

**Supplemental Table S4. Sequencing and genome mapping statistics of genomic libraries before and after CRISPR/Cas9-mediated repeat-depletion**

For each sequenced library, the table reports the type of starting samples, the type of library, the depletion protocol, the total number of sequenced fragments, the library insert size, the percentage of duplicates and the coverage statistics after normalizing to ~50 million fragments, the average mapped coverage of the nuclear genome and the fraction of nuclear sequences covered by at least 1, 2, 3, 5 and 10 reads.

| Library ID | Samples | library type | Depletion condition | Number of sequenced fragments | Mean insert size | Duplicates | Number of normalized fragments | Number of deduped mapped reads | Average mapped coverage | %1X | %2X | %3X | %5X | %10X |
|---|---|---|---|---|---|---|---|---|---|---|---|---|---|---|
| 1092 | L. culinaris cv. Castelluccio | Twist 8 plx | Not Depleted | 71,718,424 | 504.86 | 5.45% | 51,770,037 | 97,299,032 | 3.38 | 72.34 | 53.14 | 38.44 | 20.95 | 6.79 |
| 1146 | L. culinaris cv. Castelluccio | Twist 8 plx | Not Depleted | 76,186,767 | 604.05 | 5.38% | 51,770,037 | 96,650,185 | 3.34 | 74.41 | 55.73 | 40.46 | 21.28 | 5.98 |
| 4589 | L. culinaris cv. Castelluccio | Twist 8 plx | Not Depleted | 72,542,327 | 538.34 | 5.18% | 51,770,037 | 97,217,768 | 3.32 | 73.95 | 55.46 | 40.48 | 21.52 | 6.02 |
| 1145 | L. culinaris cv. Castelluccio | Twist 8 plx | All gRNA | 60,383,270 | 535.77 | 8.08% | 51,770,037 | 94,217,784 | 3.30 | 56.69 | 42.17 | 32.85 | 20.65 | 7.35 |
| 1091 | L. culinaris cv. Castelluccio | Twist 8 plx | All gRNA | 165,391,721 | 570.91 | 4.70% | 51,770,037 | 97,452,699 | 3.36 | 59.51 | 44.53 | 34.89 | 22.08 | 7.54 |
| 1306 | L. culinaris cv. Castelluccio | Twist 8 plx | All gRNA | 104,733,474 | 617.33 | 8.56% | 51,770,037 | 93,044,740 | 3.17 | 58.04 | 42.96 | 33.45 | 21.12 | 7.28 |
| 2971 | L. culinaris cv. Castelluccio | Twist 8 plx | 3 grp gRNA | 51,860,439 | 544.61 | 12.86% | 50,098,080 | 85,837,506 | 2.99 | 46.09 | 33.66 | 26.71 | 17.88 | 7.18 |
| 1144 | L. culinaris cv. Castelluccio | Twist 8 plx | 3 grp gRNA | 63,694,363 | 640.22 | 11.91% | 51,770,037 | 90,230,015 | 3.16 | 52.98 | 39.50 | 31.31 | 20.52 | 7.47 |
| 5700 | L. culinaris cv. Castelluccio | Twist 8 plx | 3 grp gRNA | 154,395,442 | 601.47 | 8.09% | 51,770,037 | 93,994,666 | 3.23 | 48.14 | 35.55 | 28.68 | 19.97 | 8.50 |
| 4017 | L. culinaris cv. Castelluccio | Twist 8 plx | 3 grp 2x gRNA | 63,592,136 | 523.03 | 9.51% | 51,770,037 | 92,705,089 | 3.19 | 45.76 | 33.67 | 27.06 | 18.66 | 8.01 |
| 1308 | L. culinaris cv. Castelluccio | Twist 8 plx | 3 grp 2x gRNA | 82,709,820 | 602.82 | 12.18% | 51,770,037 | 89,423,516 | 3.01 | 44.65 | 32.94 | 26.53 | 18.35 | 7.65 |
| 5701 | L. culinaris cv. Castelluccio | Twist 8 plx | 3 grp 2x gRNA | 71,851,660 | 561.12 | 8.04% | 51,770,037 | 93,688,924 | 3.14 | 42.64 | 31.17 | 25.51 | 18.42 | 8.50 |
| 1288 | L. culinaris cv. Redberry and L. orientalis | Twist 8 plx | Not Depleted | 73,075,884 | 610.18 | 4.93% | 51,770,037 | 97,180,421 | 3.36 | 79.00 | 59.78 | 43.33 | 22.47 | 5.86 |
| 1290 | L. culinaris cv. Redberry and L. orientalis | Twist 8 plx | Not Depleted | 114,137,748 | 627.42 | 4.47% | 51,770,037 | 97,851,366 | 3.39 | 78.24 | 58.96 | 42.82 | 22.54 | 6.18 |
| 2970 | L. culinaris cv. Redberry and L. orientalis | Twist 8 plx | 3 grp 2x gRNA | 64,897,732 | 625.67 | 10.23% | 51,770,037 | 91,661,448 | 3.15 | 44.41 | 33.21 | 27.31 | 19.78 | 9.16 |
| 4884 | L. culinaris cv. Redberry and L. orientalis | Twist 8 plx | 3 grp 2x gRNA | 104,291,293 | 593.02 | 5.20% | 51,770,037 | 96,581,457 | 3.32 | 48.39 | 36.26 | 29.61 | 21.10 | 9.44 |
| 5685 | L. culinaris cv. Redberry and L. orientalis | Twist 8 plx | 3 grp 2x gRNA | 88,665,568 | 583.77 | 6.23% | 51,770,037 | 95,285,490 | 3.23 | 46.30 | 34.34 | 28.07 | 20.15 | 9.20 |
| 5684 | L. culinaris cv. Redberry and L. orientalis | Twist 8 plx | 3 grp 2x gRNA | 57,540,234 | 578.60 | 6.61% | 51,770,037 | 95,014,227 | 3.21 | 46.03 | 34.26 | 27.99 | 20.01 | 9.06 |
| 1090 | L. culinaris cv. Castelluccio | Twist 96 plx | Not Depleted | 208,211,337 | 614.41 | 3.32% | 51,770,037 | 98,387,221 | 3.28 | 71.42 | 51.99 | 37.41 | 20.27 | 6.51 |
| 4560 | L. culinaris cv. Redberry and cv. Castelluccio | Twist 96 plx | Not Depleted | 123,998,302 | 543.78 | 4.10% | 51,770,037 | 97,483,388 | 3.17 | 72.03 | 50.94 | 35.92 | 19.53 | 6.57 |
| 4588 | L. culinaris cv. Castelluccio | Twist 96 plx | Not Depleted | 58,967,488 | 538.15 | 4.87% | 51,770,037 | 96,257,002 | 3.14 | 69.97 | 50.02 | 35.56 | 19.09 | 6.12 |
| 4014 | L. culinaris cv. Castelluccio | Twist 96 plx | 3 grp 2x gRNA | 62,479,471 | 637.44 | 7.08% | 51,770,037 | 94,921,666 | 3.18 | 37.94 | 28.20 | 23.21 | 16.93 | 8.26 |
| 4890 | L. culinaris cv. Redberry and cv. Castelluccio | Twist 96 plx | 3 grp 2x gRNA | 131,185,143 | 625.66 | 3.90% | 51,770,037 | 97,903,510 | 3.27 | 50.07 | 36.78 | 28.98 | 19.10 | 7.67 |
| 4886 | L. culinaris cv. Castelluccio | Twist 96 plx | 3 grp 2x gRNA | 127,713,553 | 638.44 | 5.39% | 51,770,037 | 96,703,552 | 3.26 | 47.46 | 34.97 | 27.89 | 18.92 | 7.94 |
| 3993 | L. culinaris cv. Castelluccio | WGS | Not Depleted | 90,835,454 | 536.14 | 3.20% | 51,770,037 | 99,666,226 | 3.86 | 81.62 | 71.40 | 58.44 | 30.94 | 3.94 |
| 6274 | L. culinaris cv. Redberry | WGS | Not Depleted | 76,123,437 | 580.96 | 7.99% | 51,770,037 | 94,698,728 | 3.66 | 90.28 | 79.35 | 62.81 | 28.29 | 1.67 |
| 6275 | L. orientalis | WGS | Not Depleted | 78,167,316 | 556.49 | 7.47% | 51,770,037 | 95,056,851 | 3.66 | 75.89 | 64.37 | 52.39 | 29.43 | 5.64 |
| 6276 | L. culinaris cv. Castelluccio | WGS | 3 grp 2x gRNA | 109,288,303 | 585.90 | 6.51% | 51,770,037 | 95,761,346 | 3.70 | 46.85 | 36.40 | 31.15 | 24.86 | 14.12 |
| 6277 | L. culinaris cv. Redberry | WGS | 3 grp 2x gRNA | 104,579,542 | 605.45 | 6.45% | 51,770,037 | 96,224,528 | 3.74 | 46.84 | 36.60 | 31.61 | 25.66 | 15.34 |
| 6278 | L. orientalis | WGS | 3 grp 2x gRNA | 107,633,126 | 573.09 | 6.40% | 51,770,037 | 95,608,177 | 3.68 | 43.46 | 33.79 | 29.07 | 23.52 | 14.41 |

**Supplemental Table S5.** ***L. culinaris*** **repeat features and efficiency of depletion**

For each repeat class present in the L. culinaris genome, the table shows the occurrence (copy number), the total length, the genomic fraction, the median lenght of each repeat unit, the total number of gRNAs targeting the repeat class and each repeat unit, the cutting frequency (gRNAs/kbp) and the variation of read mapping in that repeat type following CRISPR/Cas9-mediated repeat-depletion (bp – base pairs; FC – fold change).

| Repeat type | copy number | Total repeat length (bp) | Fraction of genome (%) | Length of repeat unit (mean) | Total gRNAs | gRNAs/Kbp | gRNAs/repeat unit (mean) | mean variation of mapped reads (log2FC) |
|---|---|---|---|---|---|---|---|---|
| LTR-Gypsy | 1728606 | 2,400,036,354 | 63.826% | 2076.64 | 35,357,754 | 14.73 | 41.07 | -0.92 |
| LTR-Copia | 629868 | 579,495,211 | 15.411% | 1163.20 | 6,404,320 | 11.05 | 14.16 | -0.04 |
| other LTR | 155019 | 127,000,455 | 3.377% | 882.02 | 1,128,270 | 8.88 | 8.04 | -0.71 |
| Line | 53111 | 37,554,666 | 0.999% | 757.98 | 150,347 | 4.00 | 3.06 | 0.50 |
| CACTA | 32589 | 23,669,215 | 0.629% | 771.26 | 114,718 | 4.85 | 3.77 | 0.35 |
| mu | 30417 | 19,464,208 | 0.518% | 761.84 | 183,711 | 9.44 | 7.70 | 0.78 |
| hAT | 10143 | 3,758,344 | 0.100% | 392.03 | 12,850 | 3.42 | 1.37 | 1.42 |
| Helitron | 6684 | 3,725,032 | 0.099% | 578.69 | 23,938 | 6.43 | 3.80 | 1.44 |
| other TE (<0.1%) | 2244 | 1,125,294 | 0.030% | 530.30 | 1,474 | 1.57 | 0.72 | 1.53 |
| Tandem Repeats (TR) | 23045 | 13,460,124 | 0.358% | 606.31 | 107,197 | 7.96 | 4.83 | 0.48 |
| Simple Repeats (SR) | na | 58,089,181 | 1.545% | 97.90 | 116,949 | 2.01 | 0.20 | 1.49 |
