## Supplemental Figures for "CRISPR/Cas9-based repeat depletion for the high-throughput genotyping of complex plant genomes"

**Supplemental Figure S1. Variation of mapped reads coverage in relation to gRNA density.** Density plot showing the variation of mapped reads coverage on nuclear repeat segments in relation to the number of gRNAs targeting the same region, in one representative experiment. Only regions with at least one gRNA cut are shown and the log2FC of coverage variation was calculated as described in the Method section. Percentages indicate the fraction of regions with more than eight gRNAs showing either positive or negative read variations. For visualization purposes, data exceeding coverage variation values of –5 or 5 and cut frequency values of 30 were plotted as points at saturation and constitute 1.6% of total dataset. FC – fold change.

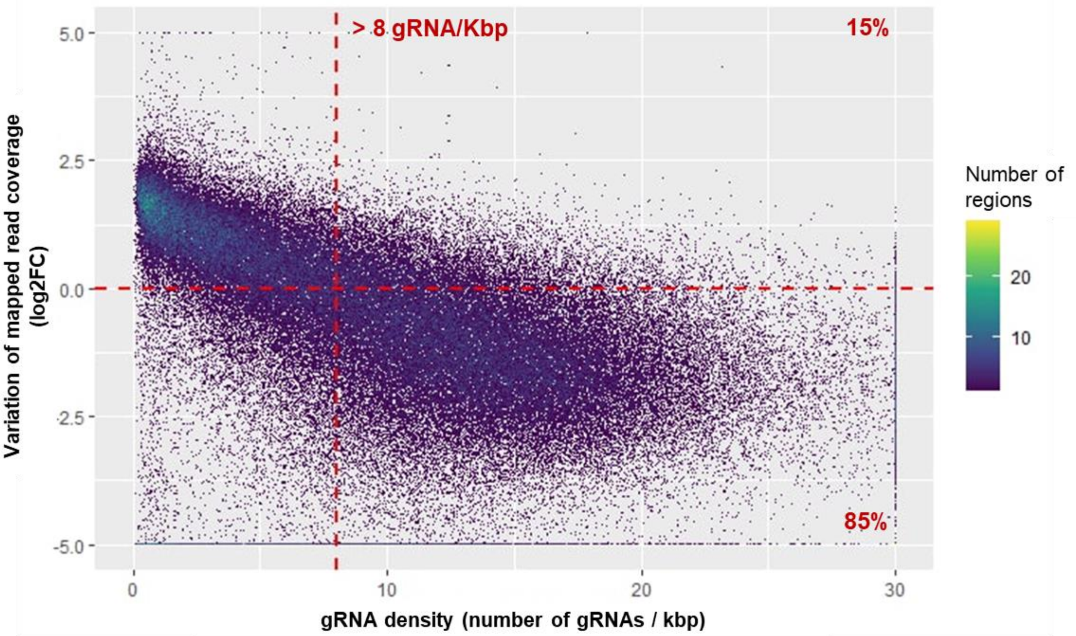

**Supplemental Figure S2. Examples of false negative variants rescued by CRISPR/Cas9-mediated repeat depletion.** Integrative Genome Browser Visualization (IGV) of Illumina sequencing reads mapped at two genomic sites of ~40 bp (**A** and **B**) before (upper tracks) and after (lower tracks) CRISPR/Cas9-mediated repeat depletion. Two heterozygous variants have been called in the depleted sample but were overlooked in the depleted sample, despite their location in a genotypable position (PASS, DP ≥ 5).

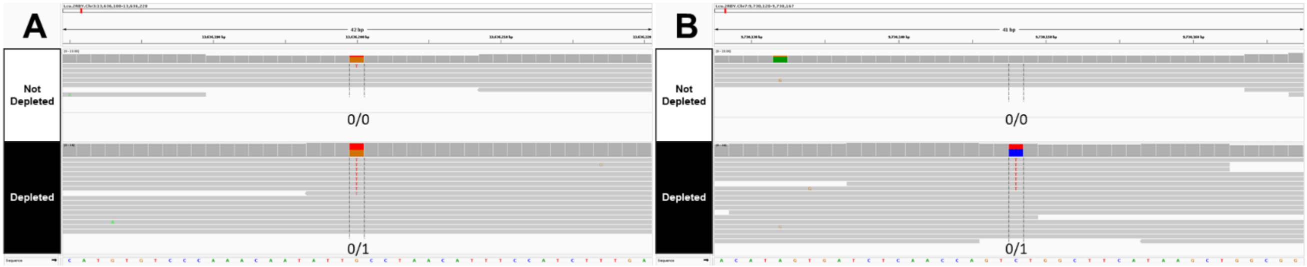
